# Germline Targeted Baboon Apolipoprotein L-1 Protects Mice Against African Trypanosomes

**DOI:** 10.1101/2025.09.23.676901

**Authors:** Sara Fresard, Sarah J. Pangburn, Kayla Leiss, Daphne Boodwa-Ko, Daniella Kovacsics, Chris J. Schoenherr, Jeremy S. Rabinowitz, Aris N. Economides, Li Li, Weigang Qiu, Bernardo Gonzalez-Baradat, Alessandro Rosa, Russell Thomson, Jayne Raper, Joseph Verdi

**Affiliations:** Biology Program, The Graduate Center at the City University of New York, New York City, NY 10016 USA; Department of Biological Sciences, Hunter College at the City University of New York, New York City, NY, 10065 USA; Regeneron Pharmaceuticals, Tarrytown, NY, 10591 USA

**Keywords:** trypanosoma brucei, apolipoprotein l1, trypanosome lytic factor, primate evolution

## Abstract

Certain primates are immune to infection by most African trypanosome parasites due to apolipoprotein L-1 (APOL1), a primate-specific ion channel-forming protein. To broaden our understanding of primate APOL1, we generated a panel of trypanosome-resistant murine models expressing various primate APOL1 proteins. We used these mice to investigate the role of APOL1 in trypanosome immunity *in vivo* by challenging them with various human and livestock trypanosome isolates. Baboon APOL1 provides partial protection to trypanosome isolates, though its protective capacity was limited by poor expression. A more highly expressed chimeric APOL1 encoding human APOL1 with the baboon APOL1 C-terminus was protective against human-infective trypanosomes, although with a fitness cost likely associated with high expression. We investigated the long-standing assumption that human resistance to *Trypanosoma vivax* is mediated by APOL1. Surprisingly, APOL1-expressing mice were fully susceptible to *T. vivax* infection, challenging this hypothesis. These model systems are useful tools for evaluating the possibility of generating genetic engineered livestock for disease control.

## Introduction

African trypanosomes proliferate in the bloodstream of mammals after transmission by the tsetse fly. Trypanosome endemicity restricts the agricultural and economic development of the African continent, resulting in 3 million cattle fatalities per year and subsequent annual losses in the range of 1-4 billion USD^1^. Human infection and mortality are less common (<1,000 reported cases per year)^2^, which can be partially attributed to efficient disease control efforts, but also to the innate immune protection afforded exclusively to primates by the haptoglobin-related protein (*HPR*) and apolipoprotein L-1 (*APOL1*) genes. HPR and APOL1 are antimicrobial proteins that circulate in sera on high-density lipoprotein (HDL) complexes called trypanosome lytic factors (TLFs)^3–5^. TLFs are endocytosed by the invading trypanosome species either by fluid phase or by receptor-mediated uptake. The latter is driven by an interaction between the trypanosome haptoglobin-hemoglobin receptor (HpHbR) and the HPR protein, driving APOL1 internalization^6^. APOL1 is a channel-forming protein that forms cation-permeable channels in parasite membranes. Channel formation by human APOL1 involves an initial membrane insertion step that requires the acidic pH of the endocytic pathway, followed by pH neutralization-mediated channel opening, which may occur after recycling to the plasma membrane^7–9^. This process ultimately leads to osmotic lysis of the trypanosome^10^.

HDLs, and therefore TLFs, are synthesized primarily in the liver before being secreted into serum^11^. HPR mRNA is produced only in the liver^12^. Currently, HPR protein has no hypothesized function outside of TLF and HDL-associated biology. Conversely, APOL1 mRNA is more ubiquitous and roles of APOL1 have been identified in intracellular viral immunity^13^ and kidney function^14^. Various kidney cell types express APOL1 without incident in the majority of the human population. However, a fraction of individuals with recent African ancestry harbor the deleterious APOL1 variants called G1 or G2 which can lead to a variety of APOL1-associated nephropathies^15^. Efforts to study these kidney-specific phenotypes *in vivo* used transgenic mice generated by driving the expression of APOL1 using kidney-specific promoters^16,17^. However, these mice do not produce detectable serum APOL1 or trypanosome lytic factors. We have previously used a model of hydrodynamics-based gene delivery (HGD) to study HPR and APOL1 function *in vivo*, in part because the main target of the plasmid injection is the liver, and we could therefore model the function of circulating TLF^18^. However, this genetic modification strategy is transient, with expression only lasting about 2 weeks. Therefore, there remains a need for a murine model that stably produces primate TLFs.

*Trypanosoma brucei rhodesiense* and *T. b. gambiense* are the only African trypanosomes that regularly infect humans. Human infection by these parasites is mediated by independently evolved APOL1 resistance mechanisms. *T. b. rhodesiense* parasites express serum resistance associated (SRA) protein, which prevents human APOL1 channels from forming by directly binding to the C-terminal domain of APOL1 and inhibiting membrane insertion^5,19,20^. However, SRA does not provide resistance against baboon (*Papio* species) APOL1. Specific amino acids near the baboon APOL1 C-terminus prevent SRA binding *in vitro*, and transient (HGD) baboon APOL1 expression protects mice from infection by *SRA*-expressing trypanosomes *in vivo*^20^. Transient expression of chimeric APOL1 proteins that encode the majority of the human APOL1 sequence fused to the baboon C-terminus also provided protection^20^. However, these experiments used transgenic laboratory isolates of *T. b. brucei* modified to overexpress the *SRA* gene, which may not be representative of clinical or field isolates. Whether these chimeric APOL1 sequences can protect against clinical isolates and/or field strains of trypanosomes remains to be tested.

The mechanism of human APOL1 resistance by *T. b. gambiense* is a combinatorial mechanism consisting of a mutated HpHb receptor, which reduces APOL1 uptake, and the *Trypanosoma gambiense-*specific glycoprotein (TgsGP) which may block APOL1 membrane insertion^21–23^. However, this resistance mechanism is not sufficient to protect against lysis by *Papio papio* APOL1^24^. The molecular mechanism by which baboon APOL1 channels bypass the function of TgsGP and lyse *T. b. gambiense* has not yet been characterized^25^. Recent work has revealed that the baboon variants of APOL1 insert into membranes and activate at a broader pH range than the human protein, which likely impacts trypanolysis efficiency^8^.

Published literature suggests differential susceptibility of various baboon species to *T. b. gambiense*. *P. papio* serum lyses *T. b. gambiense in vitro*^24^ and *P. papio* and *P. hamadryas* baboons are reported to be resistant to *T. b. gambiense* infection *in vivo*^26^. However, *P. cynocephalus* serum did not lyse *T. b. gambiense in vitro*^27^ and *P. cynocephalus* baboons are susceptible to *T. b. gambiense in vivo*^28^. Whether this differential resistance to *T. b. gambiense* infection can be attributed to the APOL1 protein remains unclear as many of these experiments were performed prior to the discovery of APOL1, and to date, only three of six baboon *APOL1* gene sequences are available (one each from *P. papio*, *P. hamadryas*, and *P. anubis*).

Here, we used whole genome sequencing data generated by the Baboon Genome Consortium^29^ to expand the number of available baboon APOL1 sequences to 33, representing all six baboon species. We then characterized the trypanolytic potential of the most common APOL1 variants specific to *P. papio*, *P. hamadryas*, and *P. cynocephalus,* revealing significant changes in protein function dependent on species-specific polymorphisms. A panel of genetically modified mouse lines were engineered to produce optimally trypanolytic baboon APOL1 variants with or without baboon HPR co-expression, providing useful resources to study the interactions between HDL-associated APOL1 and a broad range of pathogens. These experiments reveal notable differences in the dynamics of infection by a variety of trypanosome species, including *T. b. gambiense, T. b. rhodesiense, T. congolense, and T. vivax*. In particular, *T. vivax*, which is responsible for cattle infection along with *T. brucei* and *T. congolense*, is not recognized as being human infective. Despite the logical hypothesis that APOL1 plays a role, it has not been confirmed as the mechanism of human immunity^30^. We provide data that does not support this hypothesis and instead reveals that mice that express APOL1 and are protected against *T. brucei* remain susceptible to *T. vivax*. This bears particular relevance to our long-term goal of assessing whether genetically engineered APOL1-expressing livestock could provide an additional layer of disease control in Africa, the feasibility of which is discussed.

## Results

### Baboon APOL1 is polymorphic within and across baboon species

The *APOL* gene family is rapidly evolving, and specifically the *APOL1* gene displays evidence of positive selection in primates^31,32^. We investigated whether the baboon *APOL1* gene displayed similar patterns of selection. To determine whether *APOL1* polymorphisms exist within the baboon lineage, we analyzed data collected by the Baboon Genome Consortium. By analyzing data from 15 individual baboons (samples listed in Supplementary Table 1) representing all six *Papio* species, we identified 39 polymorphic sites within the 5 exons of baboon *APOL1*. These sites were used to construct haplotypes and a phylogenetic tree rooted by a closely related outgroup, the gelada baboon (*Theropithicus gelada*, baboon) (Supplementary Figure 1). Many individuals had insertions and deletions (indels) within their APOL1 sequences, which we have separately summarized for clarity at the protein level (Supplementary Table 2).

Our phylogenetic analysis revealed that the *APOL1* sequences of *P. papio* and *P. hamadryas* are more similar (Figure 1) than would be expected by their genome-wide single nucleotide variant species tree^29^. To query whether baboon APOL1 is under positive selection, we quantitated the ratio of non-synonymous polymorphisms (dN) to synonymous polymorphisms (dS) within the gene (Table 1). The dN/dS ratio (omega, ω) was estimated to be 0.5773 under a neutral model using maximum likelihood. When allowing ω to vary along the sequence, 74.9% of the sites showed ω =0, 25.1% of the sites showed ω =1, and no sites showed ω >1. While the likelihood ratio test did not statistically support the positive selection model, the single-ratio estimate of ω =0.5773 is on par with the dN/dS ratio of immune-related proteins in humans^33^.

**Figure 1.**
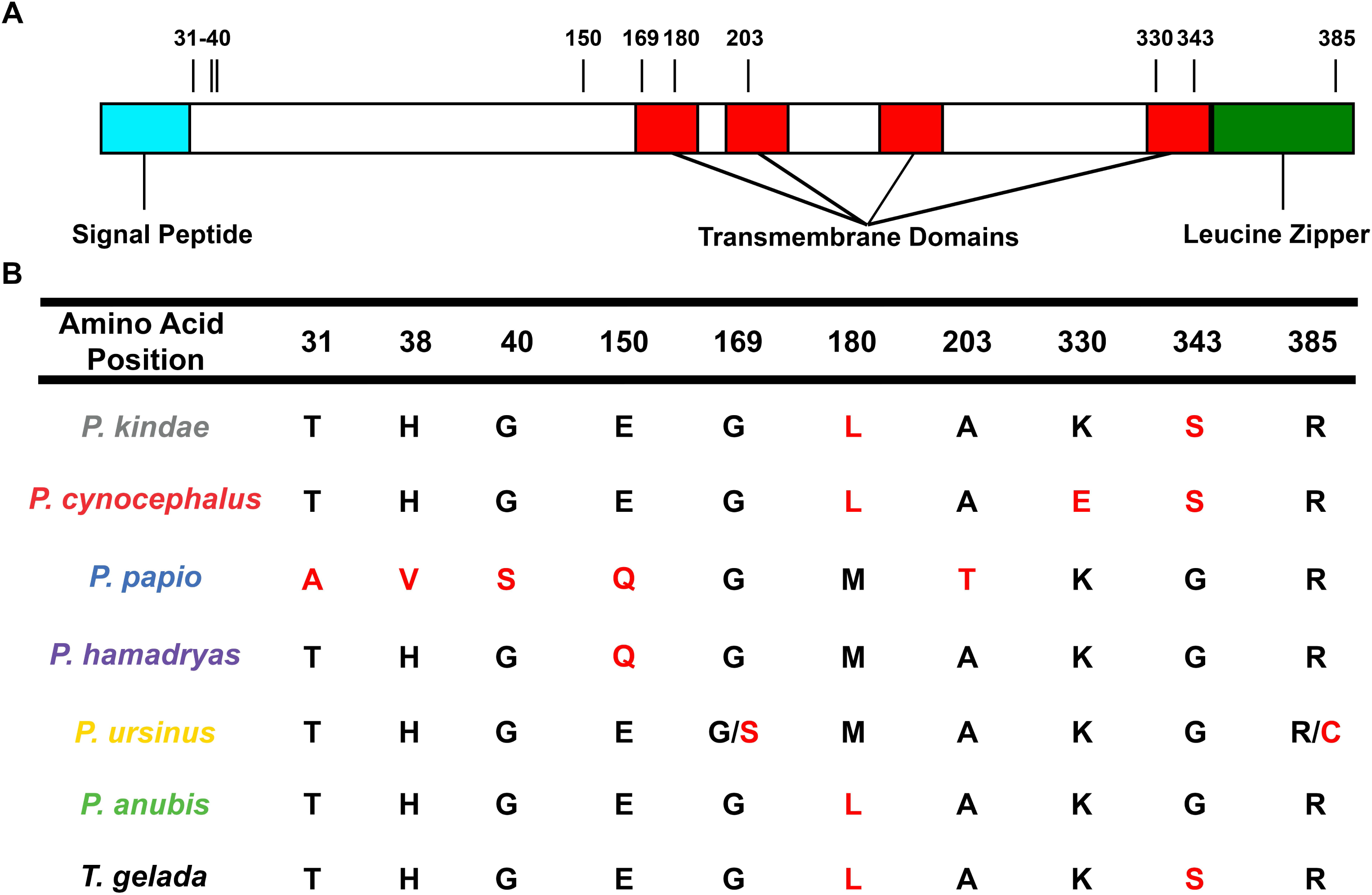
The consensus sequences of each baboon species APOL1 protein. (A) Illustration of the putative domain architecture of APOL1 (determined using JPred4), highlighting the location of the polymorphic sites that differentiate the consensus sequences of the various species. APOL1 has an N-terminal signal peptide (aqua) that directs the protein to the endoplasmic reticulum during translation, four predicted membrane spanning helices (JPred4, red), and a C-terminal leucine zipper motif (green). (B) Only the polymorphic amino acids that define the consensus sequences are listed, meaning that every non-listed amino acid position is identical between all species consensus sequences. However, more rare polymorphic sites specific to certain subgroups are listed in Supplementary Figure 1. Sites that differ from the consensus among the different species are highlighted in red. Positions 169 and 385 in *P. ursinus* APOL1 cannot be assigned given the low number of individuals (n=2), so both possible variants are listed.

**Table 1.** Analysis of synonymous (dS) and nonsynonymous (dN) substitution rates in baboon *APOL1*. The estimate from the one-ratio model (M0) (dN/dS= 0.5773) is similar to the dN/dS values of immune-related proteins from humans (28). The maximum likelihood analysis of dN/dS ratios by PAML (see Methods) indicates a nearly neutral model (M1a) where the majority (~75%) of nucleotide sites are under purifying selection with the rest of sites under neutral evolution.

| Model | kappa ($t_s/t_v$) | omega, $\omega$ ($d_N/d_s$) | lnL (likelihood) |
| --- | --- | --- | --- |
| M0 | 6.4427 | 0.5773 | -2001.0488 |
| M1a | 5.2927 | 0 (74.9%); 1(25.1%) | -1988.6290 |
| M2a | 5.2927 | 0 (74.9%); 1 (25.1%); >1 (0) | -1988.6291 |

### *P. cynocephalus* APOL1 is less trypanolytic than other *Papio* APOL1 proteins

*P. papio* APOL1 synthesized recombinantly in bacteria (rAPOL1) kills *T. b. gambiense* parasites *in vitro*^24^, and *P. hamadryas* baboons are refractory to *T. b. gambiense* infection *in vivo*^26^. *P. cynocephalus* baboons, however, are susceptible to *T. b. gambiense* infection^28^. *P. cynocephalus* serum could not kill *T. b. gambiense* parasites *in vitro*^34^, and the *P. cynocephalus APOL1* variants were divergent from the *P. hamadryas* and *P. papio APOL1* sequences in our phylogenetic analysis. To determine if the ability to kill *T. b. gambiense* was restricted to *P. papio* APOL1, we synthesized recombinant *P. papio*, *P. hamadryas, and P. cynocephalus* APOL1 variants using *E. coli* (Supplementary Figure 2A and B) and performed *in vitro* trypanosome lytic assays. Like human APOL1, all three baboon proteins killed *T. b. brucei*, however, unlike human APOL1, only the baboon proteins killed *T. b. gambiense* (Supplementary Figure 2C), suggesting that the ability to lyse *T. b. gambiense* may be afforded to all *Papio* APOL1s. In our studies, we found that *P. cynocephalus* APOL1 was less lytic to trypanosomes than *P. hamadryas* (Figure 2A), but it was still able to kill *T. b. gambiense* at higher concentrations (Supplementary Figure 2C). To determine if the concentrations required to kill trypanosomes were physiologically relevant, we quantified the circulating plasma-APOL1 concentrations in a panel of baboon samples (Figure 2B). Baboon plasma-APOL1 levels range from 100-800 ng of protein per ml of plasma (Figure 2B). The IC50s obtained in the rAPOL1 trypanosome lytic assays fell within this range of protein concentration (Figure 2A). Weak bases, such as ammonium chloride, inhibit trypanolysis by human APOL1. In contrast, we observed that while ammonium chloride treatment increased the IC50 of APOL1 from *P. hamadryas* and *P. cynocephalus*, it was not sufficient to completely block activity (Figure 2C). This suggests that the balance between *T. b. gambiense* infection and immunity in baboons is likely multifactorial, involving species-specific amino acid residues (Figure 1), differences in plasma-APOL1 levels (Figure 2B), and differences in pH regulation (Figure 2C) that are required for optimal APOL1 function.

**Figure 2.**
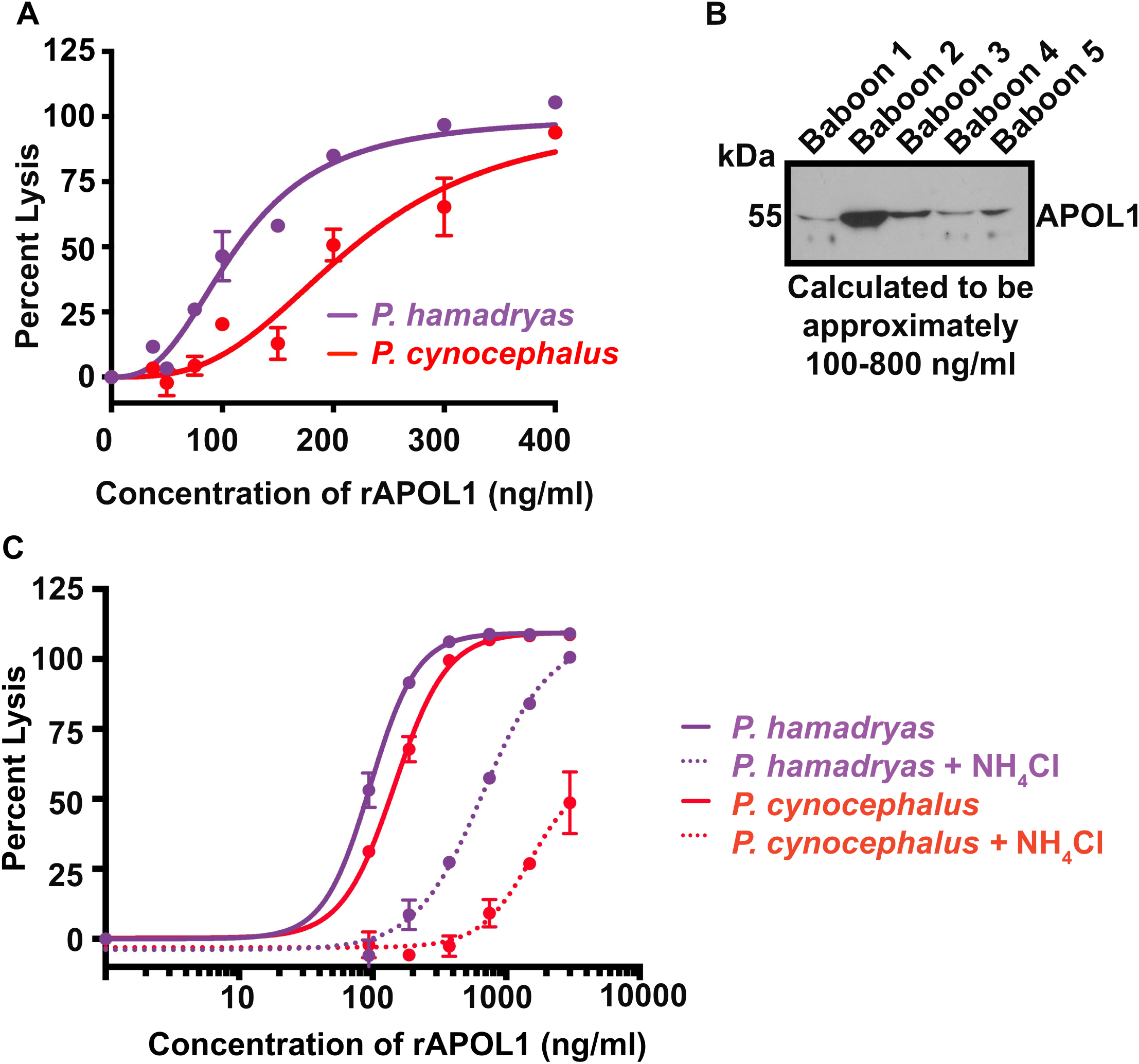
Naturally occurring baboon *APOL1* polymorphisms modulate the trypanolytic capacity of the protein. (A) 24-hour trypanolysis assay showing the lytic capacity of serially diluted *P. hamadryas* and *P. cynocephalus* rAPOL1 proteins against *T. b. brucei* parasites. IC50 *P. hamadryas* 113.7 ng/ml, *P. cynocephalus* 217.9 ng/ml. (B) Individual baboon plasma samples reveal the range of concentrations of APOL1. The band intensities compared to a serially diluted sample of rAPOL1 via pixel counts using FIJI (NIH). (C) 24-hour trypanolysis assay showing the lytic capacity of serially diluted *P. hamadryas* and *P. cynocephalus* rAPOL1 under control conditions (solid line) and when the endosomal system has been neutralized (dotted lines) using 20 mM NH4Cl treatment of the parasites 30 minutes prior to APOL1 treatment. IC50 *P. hamadryas* 96.04 ng/ml, + NH4Cl 666.4 ng/ml, *P. cynocephalus* 146.0 ng/ml, + NH4Cl 1525 ng/ml.

### *Papio* APOL1 proteins form ion channels in planar lipid bilayers

We hypothesized that the difference in lytic capacity between baboon APOL1 variants involved differences in the channel-forming mechanisms of human and *Papio* APOL1 proteins. We proceeded to characterize *P. hamadryas* and *P. cynocephalus* APOL1 in planar lipid bilayers (Figure 3A, data summarized in the paired table). Human APOL1 associates with planar lipid bilayers at pH 5.5 (Figure 3B, orange line), inserting but remaining as closed channel, and proceeds to form open cation-selective channels only when the pH is neutralized^9^ (Figure 3B, purple line), allowing for the passage of ions measured as current. In contrast, *P. hamadryas* APOL1 begins to associate with planar membranes at neutral pH^8^. Figure 3C, green line), allowing ion flow. Additional membrane insertion is stimulated by Cis acidification (Figure 3C, orange line), and the current then increases approximately three-fold upon subsequent Cis neutralization (Figure 3C, purple line). Similar to human APOL1^9^, baboon APOL1 channel conductance is maximal when the environments on both sides of the channel are neutral (Figure 3C, purple line). These data indicate that *P. hamadryas* APOL1 forms channels through a less pH-restricted process than human APOL1 and agrees with previously published results^8^. We then characterized the pH dependence of the *P. cynocephalus* APOL1 variant and observed that, similarly to *P. hamadryas* APOL1, the protein can form conductive channels after encountering an acidic pH (Figure 3D, orange line), although the protein did not permit detectable ion flux at neutral pH prior to acidification (Figure 3D, green line). To confirm these results, we analyzed the channel forming capacity of both variants on the same membrane at neutral pH and observed that only *P. hamadryas* APOL1 produced a detectable ion flux (Figure 3E). Together, this shows that both baboon APOL1s are less restricted by pH than human APOL1, as they are both able to open channels and permit cation flux at an acidic pH, unlike human APOL1. Only *P. hamadryas* APOL1, however, can insert at a neutral pH. This suggests that the increased lytic capacity of *P. hamadryas* APOL1 (Figure 2A), and the smaller shift in IC50 when subjected to neutralization (Figure 2C) could be a function of less restricted channel formation relative to *P. cynocephalus* APOL1. This likely contributes to baboon trypanosome immunity *in vivo*.

**Figure 3.**
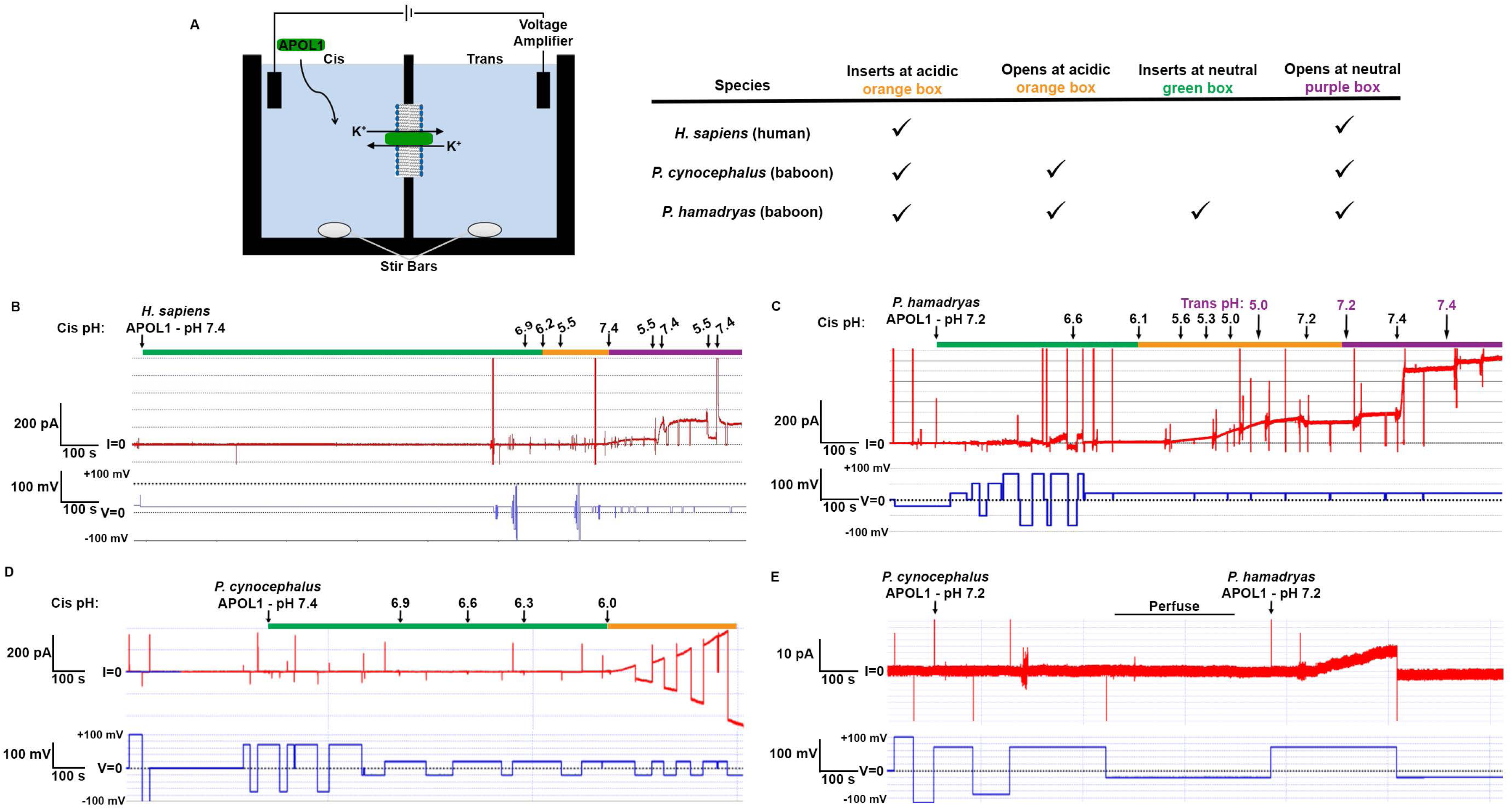
Differential channel formation by *Papio APOL1* proteins. (A) Schematic illustrating the planar lipid bilayer. Two compartments containing 1 ml buffered KCl (bilayer buffer; see methods) are separated by a phospholipid and cholesterol bilayer. The side of the bilayer that APOL1 is added to is referred to as the “Cis” side, while the opposite side is the “Trans” side. The experimenter sets the voltage (blue line, to mimic a membrane potential) and alters the pH on each side of the membrane by adding pre-calibrated volumes of KOH or HCl to the continuously stirred solutions. Formation of ion-permeant channels in the bilayer is read in the form of an electrical current (I, red line). A table summarizing the pHs at which each protein can insert and open is on the right. (B-E) The voltage (in millivolts) is shown in blue in the lower traces, and the current output (in picoamperes, pA) is shown in red in the upper traces. The green boxes indicate neutral pH, prior to acidification. The orange boxes show acidification. The purple boxes indicate neutralization after acidification. (B) The channel forming capacity of *H. sapiens*. (C) The channel forming capacity of *P. hamadryas* APOL1 at Cis (black) and Trans (purple) pH values. Ion fluxes (20-50 pA) are detected at pH 7.2 prior to Cis acidification (green box). (D) The channel forming capacity of *P. cynocephalus* APOL1 at different Cis pH values. In contrast to (B) *P. hamadryas*, there is no detectable ion flux in this region (pH 7.4-6.3, green box) prior to Cis acidification (pH 6.0, orange box), whereupon there is increasing ion flux. (E) The channel forming capacities of *P. cynocephalus* APOL1 and *P. hamadryas* APOL1 at neutral pH on the same bilayer. During the period marked with the black bar (labelled as ‘Perfuse’) the cis compartment was perfused with chamber buffer (pH 7.2) to remove non-membrane-associated protein.

### Integration of the genomic baboon *APOL1* locus into the mouse genome does not protect from trypanosome infection

Baboon and human *APOL1* are predicted to encode a 42 kDa protein. However, baboon APOL1 migrates at 55 kDa on SDS-PAGE gels, suggesting that it undergoes post-translational modification. Peptide *N*-glycosidase F (PNGase F) treatment of baboon HDL resulted in a gel mobility shift in APOL1, revealing that the protein is *N*-glycosylated (Supplementary Figure 3). However, de-glycosylation does not produce the expected 42 kDa human APOL1-sized protein, suggesting that other post-translational modifications are present. Importantly, all previous reports directly showing that *Papio* APOL1 proteins are capable of lysing trypanosomes have used sub-physiological levels of *Papio* serum or recombinant APOL1 isolated from *E. coli*, which lack the enzymes required for the addition of N-linked glycans^35^.

In order to investigate the function of baboon APOL1 in a physiologically relevant manner, we generated a panel of genetically modified mice expressing baboon *APOL1* utilizing different combinations of promoters and constructs. To build our first construct, we obtained a 20 kilobase segment of the *P. anubis* genome in a bacterial artificial chromosome (BAC) from the Children’s Hospital Oakland Research Institute. The BAC contained all five *APOL1* exons, the putative endogenous promoter, and the upstream and downstream untranslated regions of the gene which were predicted to contain the endogenous regulatory regions required for expression. This *P. anubis* DNA was isolated from the individual animal used for the baboon reference genome Panu2.0, which differs from the *P. anubis APOL1* consensus sequence at two positions. We inserted this *P. anubis APOL1* construct into the *ROSA26* locus (Supplementary Figure 4A), a ubiquitously euchromatic region of the genome commonly used for transgene placement in mice. In another construct, we generated a version of the BAC that encoded a *P. hamadryas APOL1* gene by inserting two separate base pair changes that encoded the E150Q and L180M substitutions (Figure 1). The *P. hamadryas APOL1* construct was inserted into the mouse genome immediately upstream of the *myosin heavy chain 9* (*Myh9*) gene (Supplementary Figure 4B), which is next to the endogenous location of *APOL1* in primates. We tested whether the germline transgenic mice could express and secrete APOL1, and could not detect plasma or HDL-associated APOL1. We detected *APOL1* RNA production in liver lysates (Supplementary Figure 4C), suggesting that the loci were transcriptionally active. The mice were not protected from infection by *T. b. brucei* parasites (Supplementary Figure 4D). This was consistent with previously published results using transiently transgenic animals in which baboon *APOL1* expression could only clear a trypanosome infection in mice that also expressed human *APOA-I*^36,37^. Therefore, we crossed our transgenic *APOL1*-expressing mice with human *APOA-I*-expressing mice, to potentially generate HDLs carrying both human APOA-I and baboon APOL1. However, these double transgenic mice were also susceptible to infection by *T. b. brucei* (Supplementary Figure 4E). Co-expressing baboon *HPR* by HGD in the *P. anubis APOL1*-expressing mice (Supplementary 5A) was able to provide a slight but not statistically significant level of protection from *T. b. brucei* infection (Supplementary Figure 4F). This suggests that although we were unable to detect any APOL1 protein in plasma or HDL, there may be a small amount of circulating and functional APOL1 protein in the sera of these mice stabilized by HPR. Nevertheless, we concluded that this genetic modification strategy did not produce a suitable model system.

### Stable expression of the *P. hamadryas APOL1* cDNA protects mice from trypanosome infection

We next proceeded to target the *ROSA26* locus with a more controlled *APOL1* expression system by integrating the cDNA of *P. hamadryas APOL1* under the control of a mouse-compatible ubiquitin promoter (Figure 4A) to attempt to generate transgene expression in a physiological manner, given that *APOL1* expression is ubiquitous^12^. We used cDNA to reduce the size and complexity of the construct for improved cloning efficiency. These mice produced serum circulating APOL1 protein, albeit less than an average baboon, with the homozygous mice producing approximately twice as much protein as heterozygous mice (Figure 4B). Importantly, the APOL1 in the mouse plasma and baboon plasma are the same molecular weight, suggesting that the mice post-translationally modify the protein physiologically (Figure 4B). The APOL1 was present on HDL isolated from mice, thus forming a TLF-like complex (Figure 4C). To test whether the mice were protected from *T. b. brucei* infection, we infected the mice with 5000 *T. b. brucei* parasites. An inoculum of 5000 parasites is in the range of parasites transmitted by an individual tsetse bite, although the range is very large and not well defined (0-40,000 cells per bite; mean of 3,200)^38^. With this inoculum of parasites, the *APOL1*-expressing mice were fully protected from infection (Figure 4D). Notably, this protection was achieved in mice expressing murine *APOA-I*, contrary to the previously published data using transiently transgenic mice that required human *APOA-I*^36,37^.

**Figure 4.**
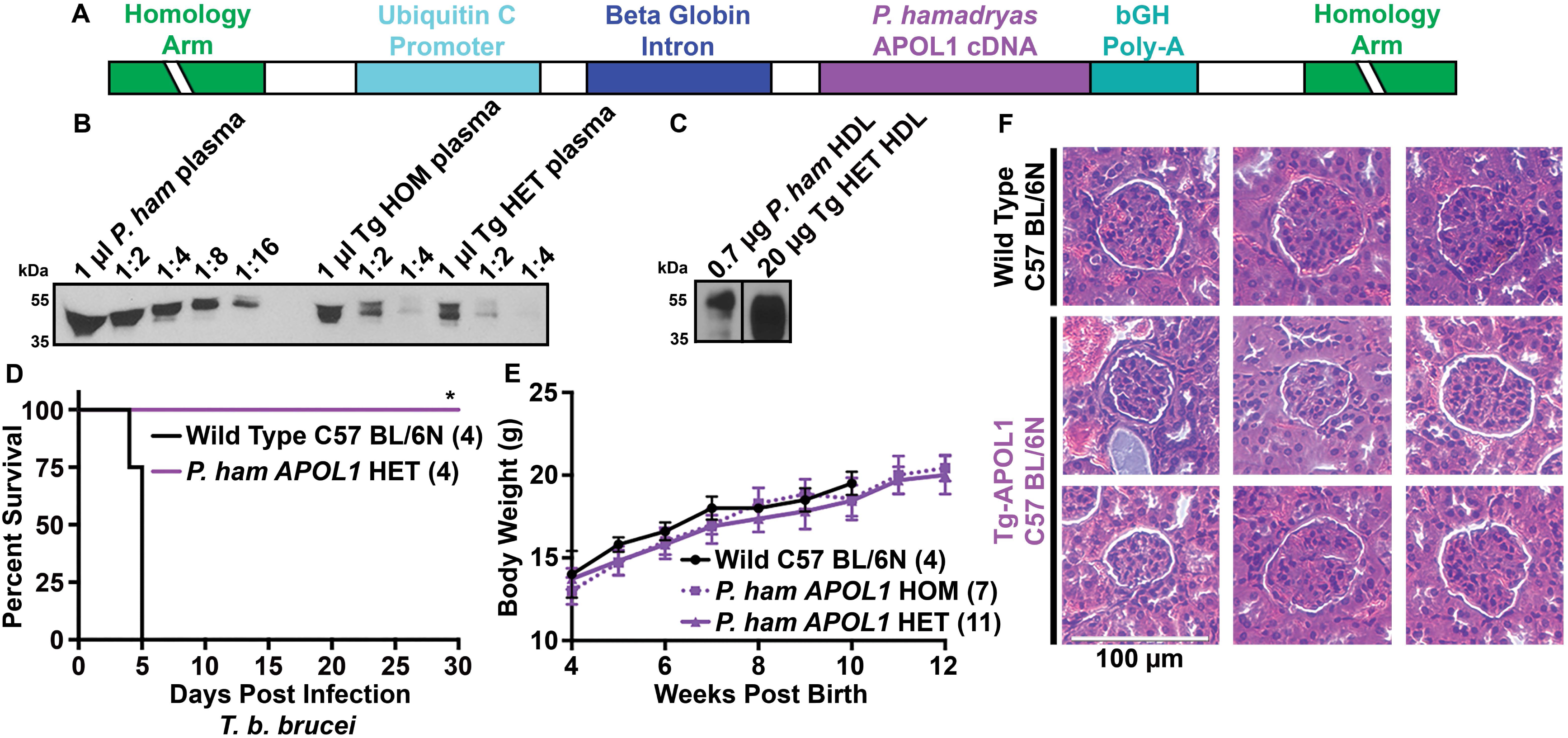
Targeted integration of the *P. hamadryas APOL1* cDNA driven by a ubiquitin promoter protects from *T. b. brucei*. (A) Schematic of the targeting construct used to insert a single copy of the *P. hamadryas APOL1* cDNA into the *ROSA26* locus of the mouse genome. bGH is bovine growth hormone. (B) Anti-baboon APOL1 western blot of serially diluted baboon *P. hamadryas* (*P. ham*) plasma and transgenic (Tg) homozygous (HOM) and heterozygous (HET) mouse plasma samples. (C) Anti-baboon APOL1 western blot of purified baboon HDL and transgenic mouse HDL samples. (D) Kaplan-Meier curve showing the survival of *P. hamadryas APOL1* transgenic heterozygous mice compared to wild type counterparts infected with 5000 *T. b. brucei* Lister-427 parasites i.p. (* p = 0.01; Log-rank test). (E) Weight gain in heterozygous and homozygous APOL1 transgenic mice compared to wild type counterparts as a function of time post-birth (error bars represent the mean +/− the SD for each group each week). (F) Hematoxylin and eosin-stained kidney tissue slices focused on a glomerulus from adult wild type and *APOL1* HET transgenic mice. Images represent three mice from each group. Images were captured using a light microscope with 40X magnification.

We next assessed general animal welfare of the transgenic lines. We used multiple litters of mice to test whether the *APOL1* expression had any apparent negative impact on early post-birth development. We observed that the mice gained weight comparable to wild type counterparts (Figure 4E) and exhibited normal behavior. In humans, the G1 and G2 APOL1 variants can cause chronic kidney diseases in adults, a common manifestation of which is focal segmental glomerulosclerosis (FSGS)^14^. As FSGS progresses, the glomeruli of the kidney begin to decay over time as progressive glomerular cell death leads to the formation of scar tissue. Histologically, the glomeruli resect from the walls of the Bowman’s capsule, losing their approximately spherical shape. Importantly, glomerular damage has also been observed in mice expressing the nephrotoxic human APOL1 variants, using either kidney-specific promoters^16^ or the entire genomic region of APOL1^39^. We therefore prepared kidney tissue sections from adult (greater than 1 year old) transgenic baboon APOL1 and wild type mice for basic histological analyses and observed that the glomeruli in the transgenic mice showed no obvious signs of resection and instead resembled wild-type glomeruli (Figure 4F). We concluded that in this mouse line, baboon APOL1 did not induce any detectable negative outcomes in the tested phenotypes.

Once we determined that baboon APOL1 expression caused no obvious deleterious effects, and provided protection from infection by *T. b. brucei*, we proceeded to characterize the protective potential of *P. hamadryas APOL1* expression in the context of a wide range of human-infective and livestock-infective African trypanosomes. Similar to *T. b. brucei* (Figure 4D), the *APOL1*-expressing homozygote mice were fully protected from infection by the globally distributed livestock pathogen *T. b. evansi* (Table 2). However, the mice were only partially protected from human-infective *T. b. gambiense* (Table 2). After infecting mice with 5000 *T. b. gambiense* parasites, we found that homozygous mice were partially protected from infection (Table 2). Inoculating homozygous mice with only 500 parasites extended the survival of the mice significantly, although they still were not fully protected (Table 2). We then investigated whether the mice were protected from infection by *T. b. rhodesiense*, a zoonotic pathogen that readily infects both humans and cattle. Heterozygous mice infected with *T. b. rhodesiense* were partially protected, with a survival extension of approximately 2 to 3 weeks relative to wild-type controls (Table 2). Homozygous *APOL1*-expressing mice were fully protected from infection (Table 2), suggesting that *APOL1* levels are an important determining factor that dictates whether an animal will be fully resistant to infection. We were also able to achieve nearly complete protection by co-expression of *HPR* via HGD in the heterozygous mice (Table 2), affirming that TLFs (containing APOL1, HPR, APOA-I) that can both kill the parasites and be endocytosed by the HpHbR are more effective than TLFs (containing APOL1, APOA-Il; lacking HPR) that must be taken up by fluid-phase endocytosis.

**Table 2.** *P. hamadryas* APOL1 germline transgenic mice are partially protected from infection by *T. brucei* subspecies. Mice (transgenic; purple, or wildtype; black) were inoculated with the indicated subspecies of trypanosome and survival was monitored. The mice infected with *T. b. rhodesiense* post-hydrodynamic gene delivery (HGD) of HPR were infected 2 days after plasmid injection. Data are expressed as the number of surviving mice/number of inoculated mice. The loss of mice due to high parasitemia (day 20-50) indicates the evolution of parasites resistant to *P. hamadryas* APOL1.

| Parasite (Inoculum) | Mouse Genotype | Alive/Inoculated |  |  |  |  |
| --- | --- | --- | --- | --- | --- | --- |
|  |  | Day | 5 | 20 | 35 | 50 |
| *T. b. evansi* | *APOL1* HOM |  |  |  |  | 4/4 |
| AnTat 3 (5000) | Wild type |  |  |  |  | 0/4 |
| *T. b. gambiense* | *APOL1* HOM |  | 5/5 | 0/5 |  |  |
| ELIANE (5000) | Wild type |  | 0/5 | 0/5 |  |  |
| *T. b. gambiense* | *APOL1* HOM |  | 5/5 | 3/5 | 3/5 | 2/5 |
| ELIANE (500) | Wild type |  | 5/5 | 0/5 | 0/5 | 0/5 |
| *T. b. rhodesiense* | *APOL1* HOM |  |  | 5/5 | 5/5 | 5/5 |
| KETRI 243 (5000) | *APOL1* HET |  |  | 5/6 | 0/6 | 0/6 |
|  | *APOL1* HET + *HPR* HGD |  |  | 4/4 | 4/4 | 4/4 |
|  | Wild type + *HPR* HGD |  |  | 0/6 | 0/6 | 0/6 |
|  | Wild type |  |  | 0/9 | 0/9 | 0/9 |

We then investigated whether the mice were protected from infection by the livestock infective *T. congolense*. This strain has a variable effect on wild-type mice, causing chronic infections rather than acute. Therefore, instead of monitoring survival, we periodically measured parasitemia in the mice (Table 3). One individual heterozygote *APOL1*-expressing mouse developed a high parasitemia and subsequently died within the first 2 weeks of infection, although this was an isolated result that was never replicated (Table 3, row 2, “1/13”). More reproducibly, we observed that approximately 25% of the heterozygous *APOL1*-expressing mice developed a high and sustained parasitemia approximately 5-6 weeks after infection (Table 3, row 2, “4/13”). This was never observed in homozygous *APOL1*-expressing mice or *APOL1* and *HPR*-expressing mice (Table 3, row 1 and row 3), consistent with previous data using *T. b. rhodesiense* (Table 2).

**Table 3.** *P. hamadryas* APOL1 germline transgenic mice are partially protected from infection by *T. congolense.* Mice were inoculated with 5000 parasites of each indicated strain of *T. congolense* and parasitemia was monitored by light microscopy at least every 3 days. Mice were infected 2 days after plasmid injection by HGD. *T. congolense* STIB 68-Q-APOL1R was derived from a heterozygote mouse infected with *T. congolense* STIB 68-Q that developed a sustained parasitemia resistant to APOL1. The STIB 68-Q-APOL1-revert strain was derived from a STIB 68-Q-APOL1R parasite population that was passed through a wild type mouse (in the absence of APOL1) for one month. Due to the nonlethal *T. congolense* parasitemia, data are represented as the number of mice with detectable parasites/number of mice inoculated. The growth of parasites within transgenic mice indicates the evolution of parasites resistant to *P. hamadryas* APOL1.

| Parasite | Mouse Genotype | Detectable Parasites/Inoculated |  |  |  |
| --- | --- | --- | --- | --- | --- |
|  |  | Day | 15 | 30 | 45 |
| *T. congolense*<br>STIB 68-Q | *APOL1* HOM |  | 0/5 | 0/5 | 0/5 |
|  | *APOL1* HET |  | 1/13 | 1/13 | 4/13 |
|  | *APOL1* HET + *HPR* HGD |  | 0/4 | 0/4 | 0/4 |
|  | Wild type |  | 7/7 | 7/7 | 7/7 |
| *T. congolense*<br>STIB 68-Q-APOL1R | *APOL1* HOM |  | 4/5 |  |  |
|  | *APOL1* HET |  | 2/2 |  |  |
|  | Wild type |  | 6/6 |  |  |
| *T. congolense*<br>STIB 68-Q-APOL1R-revert | *APOL1* HOM |  | 0/3 | 3/3 |  |
|  | Wild type |  | 3/3 | 3/3 |  |

### Previously APOL1-susceptible trypanosomes can develop APOL1 resistance in animals with sub-lethal levels of APOL1

The *T. b. gambiense*, *T. b. rhodesiense*, and *T. congolense* parasites that grow and survive in the transgenic mice are under continuous pressure from the baboon TLF and therefore represent parasites that have acquired APOL1 resistance. We investigated this phenotype using *T. congolense* by harvesting parasites from a heterozygote mouse that developed a sustained detectable level of parasitemia after an initial infection, and re-infecting those parasites (*T. congolense* STIB 68-Q-APOL1R) into naïve *APOL1*-expressing mice. We observed that the heterozygous and the majority (80%) of the homozygous *APOL1*-expressing mice infected with this derived strain of *T. congolense* developed a high parasitemia within the first two weeks of infection (Table 3, row 5 and 6). We therefore concluded that the parasites had indeed developed APOL1 resistance, either through selection of pre-existing minor populations of resistant parasites or *de novo* mutation(s).

APOL1-resistant *T. congolense* displayed a slowed growth phenotype (reaching peak parasitemia in 8-12 days) compared to wild-type parasites (5 days). We hypothesized that the APOL1 resistance was associated with a fitness cost, and parasites would therefore lose resistance in the absence of the selection pressure mediated by APOL1. To investigate this, we infected wild-type mice with the resistant parasites (*T. congolense* STIB 68-Q-APOL1r). We harvested the parasites one-month post-infection, hypothetically long enough for the resistant population to be outgrown by APOL1-susceptible revertants. These parasites (*T. congolense* STIB 68-Q-APOL1R-revert) were then re-infected into wild type mice and homozygous *APOL1* mice, revealing that homozygous mice were once again resistant to the infection, albeit only partially, with approximately a two-week delay in the onset of detectable parasitemia relative to wild-type mice (Table 3). We have not determined the genetic source of the resistance, although this revertible phenotype is reminiscent of the SRA-mediated APOL1 resistance mechanism acquired by *T. b. rhodesiense* which is also readily ‘turned off’ in the absence of APOL1 selective pressure^40^.

We then investigated where resistant parasites may have emerged in the APOL1 heterozygous mice. HDLs circulate in serum and permeate extravascular tissue spaces. The concentration of HDL in those tissue spaces is approximately 10-fold lower than the concentration that circulates in serum^41^. Since *T. b. brucei* invades and adapts to extravascular tissue spaces^42^, we hypothesized that APOL1-resistant *T. congolense* clones could emerge from tissue-resident trypanosomes exposed to this sub-lethal level of TLF. We observed that during the early stage of infection wild-type mice, there were a relatively small but detectable number of parasites in the tissues, and that the proportion of tissue-resident parasites increased as the infection progressed (Figure 5A). Of the analyzed tissues, we observed that the adipose tissue and the lung were the most frequently enriched tissues (Figure 5A), suggesting that the parasites may have particular tropism for different tissues. Enrichment in both tissues was more evident at the later stage of the infection (17 dpi).

**Figure 5.**
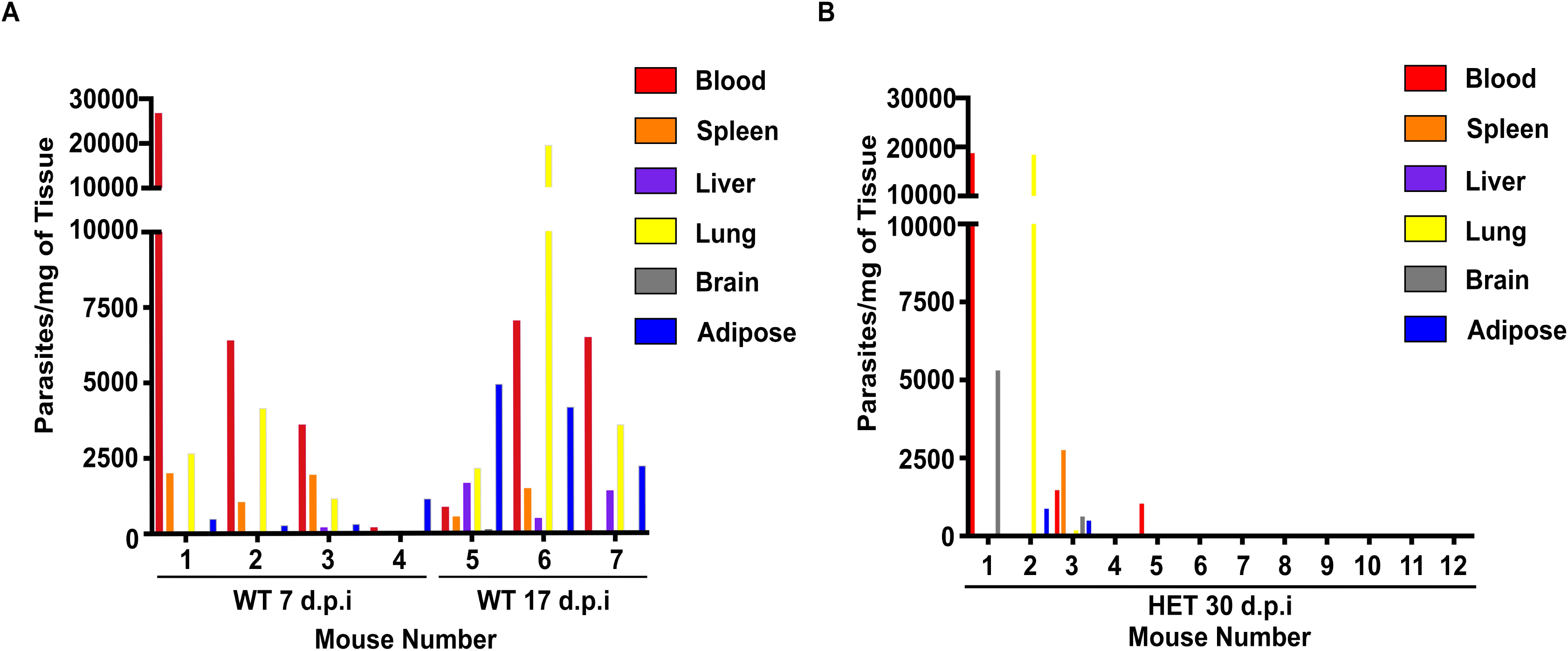
Detection of *T. congolense* DNA in tissues. (A-B) The calculated number of parasites per milligram of perfused (all blood removed) tissue in wild type (WT) C57 BL/6N-J mice (A) or heterozygous (HET) *P. hamadryas APOL1*-transgenic mice (B) after DNA isolation and qPCR using primers specific to *T. congolense Cathepsin L* (housekeeping gene) compared to a standard curve of a known number of parasites. Each number denotes 1 individual mouse. D.P.I.; Days Post-Infection.

We then investigated whether there were tissue-resident parasites in *APOL1*-expressing mice. We infected 12 heterozygous mice and harvested tissues at 30 days post-infection, hypothetically immediately preceding parasite emergence into the blood. We detected parasites in four of the mice, with one mouse (Mouse 2) displaying parasites exclusively in the lung and adipose tissue, with no detectable blood-resident parasites (Figure 5B). Notably, no parasites were ever detected in homozygous *APOL1*-expressing mice. These data suggest that tissues could provide a niche with sub-lethal levels of APOL1 that selects for the emergence of resistance.

### Co-expression of *HPR* and *APOL1* augments protection against trypanosome infection

The data thus far showed that co-expression of APOL1 (stable, germline) and HPR (HGD, transient) provided better protection than APOL1 alone. We next generated mice that stably co-express *APOL1* and *HPR* in the germline. The *P. hamadryas HPR* cDNA was expressed via an albumin promoter to drive liver-specific expression of the transgene, while *APOL1* was maintained under the same ubiquitin promoter that we previously used (Figure 6A). These mice produced similar APOL1 levels compared to our previous line of mice, and they produced HPR protein at a concentration that was comparable to baboons (Figure 6B). Heterozygote *APOL1/HPR* mice were fully protected from infection against a *T. b. brucei* strain that can infect mice that only express *APOL1* (Figure 6C). However, the protection against infection by *T. b. rhodesiense* was similar to the mice expressing only *APOL1* (Figure 6D). We also generated mice that co-expressed *APOL1* and *HPR* on separate ubiquitin promoters, anticipating that simultaneous expression of both genes would facilitate protein co-secretion onto HDLs (Figure 6E, F). These heterozygote mice were only partially protected against *T. b. brucei*, but fully protected against *T. b. rhodesiense* (Figure 6G, H). This is the first germline transgenic mouse model to provide complete protection against *T. b. rhodesiense*. With *T. b. brucei*, there was the same trend of homozygote mice being protected while some heterozygote remained susceptible. We therefore conclude that this model still has insufficient concentration of circulating APOL1 to provide complete protection against all relevant trypanosome species.

**Figure 6.**
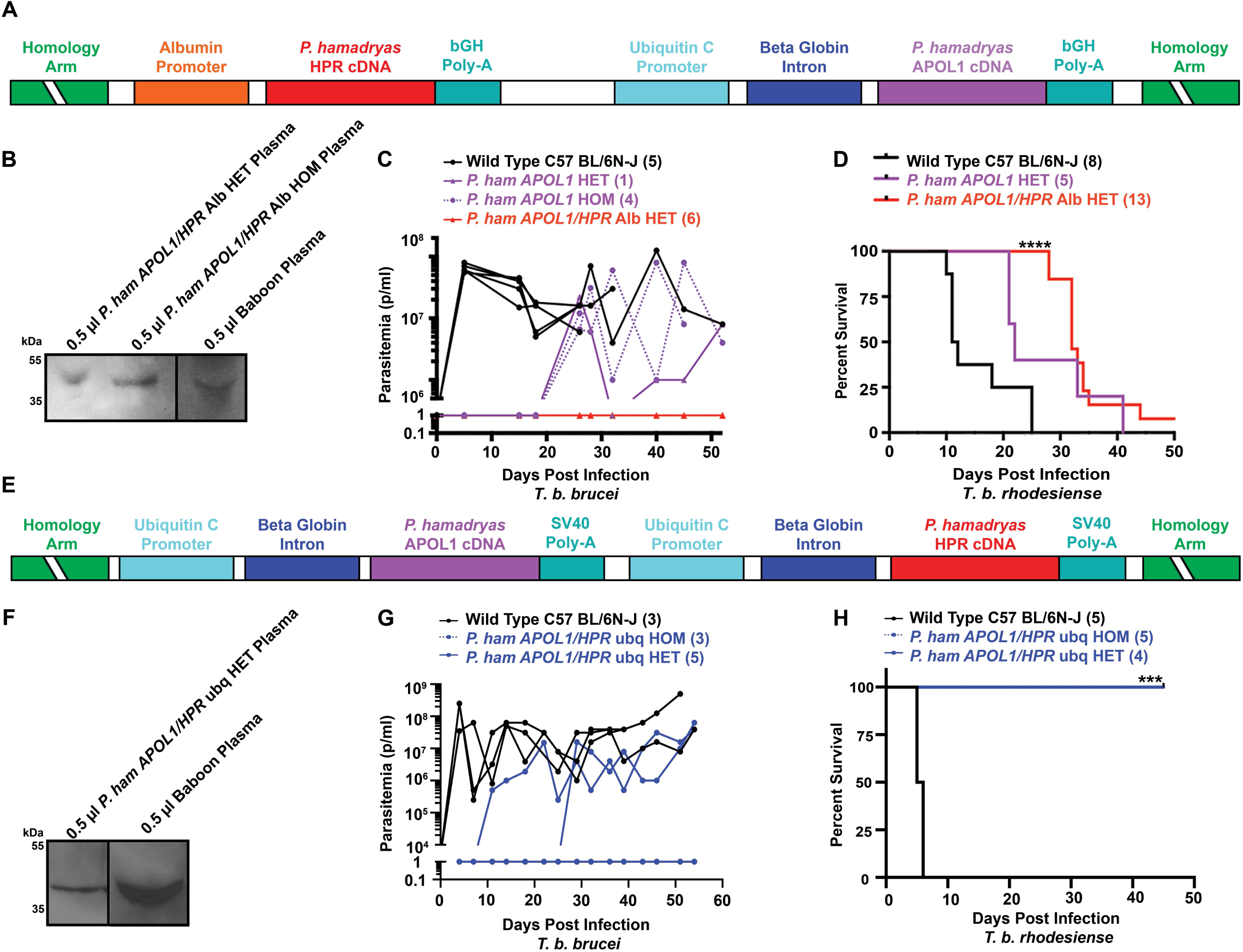
Targeted integration of *P. hamadryas APOL1* and *HPR* is not sufficient to mediate full protection against trypanosome infections. (A) Schematic of the targeting construct used to insert the *P. hamadryas APOL1* and *HPR* cDNAs into the *ROSA26* locus of the mouse genome with *APOL1* expression driven by a ubiquitin promoter and *HPR* by an albumin (alb) promoter. bGH is bovine growth hormone. (B) Western blot showing the relative concentration of HPR in the plasma of heterozygous (HET) and homozygous (HOM) transgenic mice compared to a representative baboon plasma sample using an anti-human Hp antibody that cross-reacts with human and primate HPR. (C) Parasitemia over time showing the number of parasites per mL of blood in transgenic mice inoculated with 5000 pleomorphic *T. b. brucei* (AnTat DRF) parasites by i.p. on day 0. The graph shows the parasitemia in each individual mouse of each genotype (number of mice per genotype, including wild type, in parentheses). Parasitemia was counted by light microscopy; the limit of detection is approximately 3-5 x 10^6^ parasites per mL (p/mL). The *P. hamadryas* APOL1/HPR Alb HET cohort are represented schematically as 1, though we did not detect parasites in these mice. (D) Kaplan-Meier survival curve of wild-type and HET transgenic mice inoculated with 5000 *T. b. rhodesiense* (KETRI 243) parasites i.p. (**** p < 0.0001; Log-rank test). (E) Schematic of the targeting construct used to insert the *P. hamadryas APOL1* and *HPR* cDNAs into the *ROSA26* locus of the mouse genome with *APOL1* and *HPR* expression both driven by a ubiquitin (ubq) promoter. (F) Western blot showing relative APOL1 concentration in plasma of HET transgenic mice compared to a representative baboon plasma sample, using an APOL1 antibody specific for baboon APOL1. (F) *T. b. brucei* (AnTat DRF) parasite challenge in mice WT, HET and HOM for *APOL1* and *HPR* both on ubiquitin promoters in the same scheme as outlined in C. (G) Kaplan-Meier survival curve of WT, HET, and HOM infected with *T. b. rhodesiense* (KETRI 243) in the same scheme as (D) (*** p = 0.0010; Log-rank test).

### APOL1 chimeras that do not interact with SRA, but are expressed at higher levels than the baboon protein, fully protect mice from trypanosome infection

Since co-expressing *HPR* was not sufficiently protective by this genetic modification strategy, we considered increasing the levels of circulating APOL1 to generate fully protected mice. Human APOL1 expressed in mice by HGD using similar constructs presented here circulates at about the same concentration as APOL1 in humans^43–45^, while baboon APOL1 expressed in mice (germline) circulates at a slightly lower concentration than in baboons (Figure 4B). Given that both of these models use the same expression system, we reasoned that something intrinsic to the protein coding region of the gene dictates the differences in the amount of protein detectable in serum. In order to generate mice that produce APOL1 levels high enough to protect from the important veterinary trypanosomes, we decided to produce *APOL1* chimeras encoding the majority of either the human or the closely-related gorilla *APOL1* coding sequence fused to the C-terminus of the baboon *APOL1* (human or gorilla amino acids up to and including residue 360, followed by baboon residues until the C terminus) (Hum360). The minimum-required portion of the C-terminus for ideal trypanolytic function was chosen based on the interactions of the C-terminus and the SRA protein^20,37^. The gorilla sequence was synthesized to mitigate any potential future complications related to using human DNA in a genetically modified organism (Gor360) (Figure 7A). Both chimeras were expressed at levels equivalent to human APOL1 when expressed in mice by HGD (Supplementary Figure 5A), and both chimeras fully protected mice from infection by a clinical isolate of *T. b. rhodesiense* (Figure 7B). We moved forward with these constructs and generated mice expressing either the *Hum360 APOL1* or *Gor360 APOL1* as stable transgenic animals on ubiquitin promoters. Heterozygote transgenic *Hum360* mice were protected against *T. b. rhodesiense* (Figure 7C) However, we have not been able to generate *Hum360* homozygote animals, and *Gor360* males were never able to sire. These experiments have provided the first two APOL1-only constructs capable of fully protecting mice from *T. b. rhodesiense* through genetic modification: the chimeric Hum360 (transient and stable) and the Gor360 (transient only). These constructs illustrate that high APOL1 serum protein levels are one strategy by which full trypanosome protection can be afforded, provided that toxicity is mitigated.

**Figure 7.**
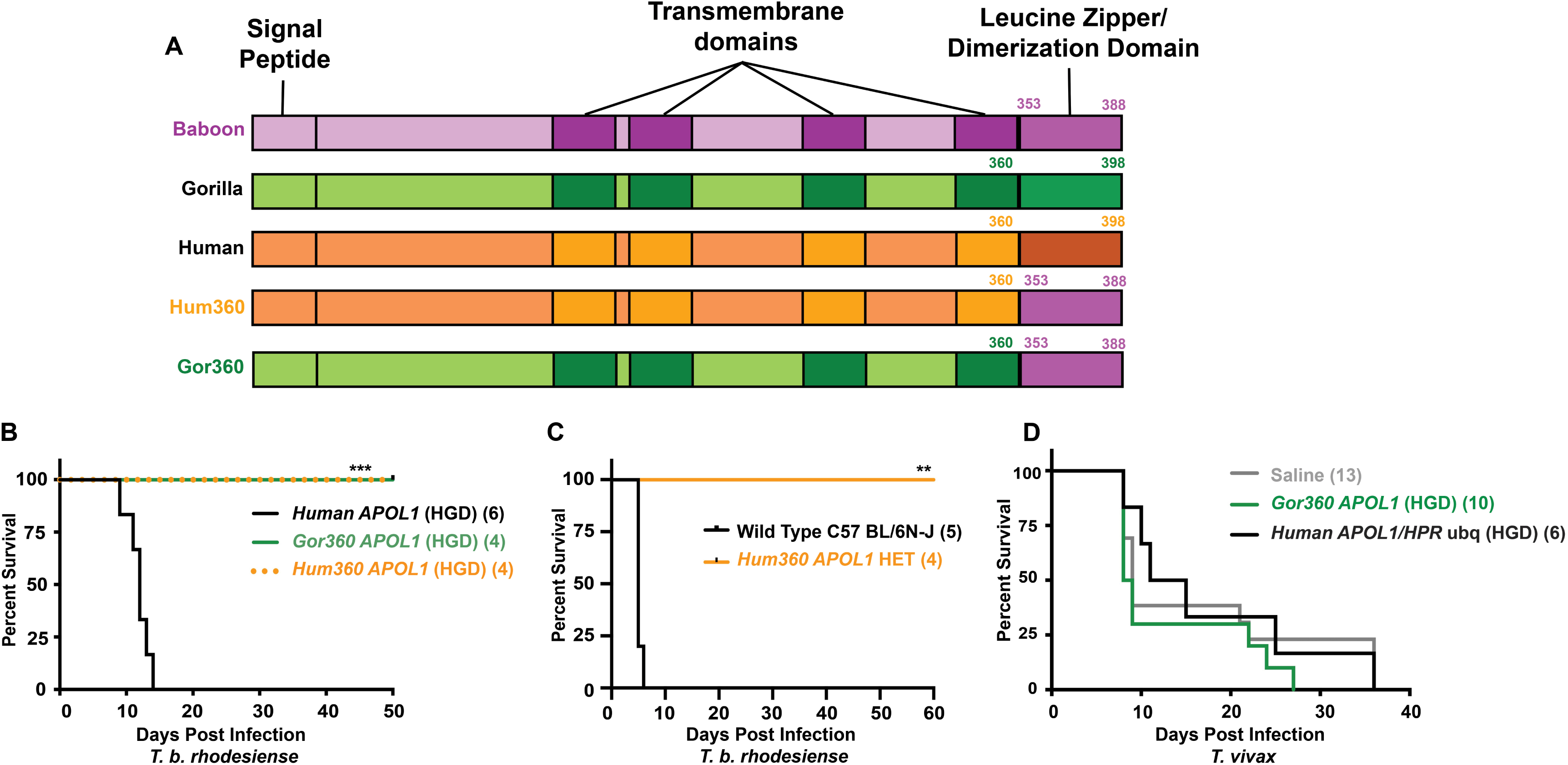
Transient HGD expression of APOL1 chimeras fully protect mice from trypanosome infection. (A) Illustration of the putative domain architecture of APOL1 (determined using JPred4 [9]) from human APOL1, gorilla APOL1, baboon APOL1, and the chimeric *APOL1* genes, which encode the first 360 amino acids of either the human (*Hum360*) or gorilla (*Gor360*) APOL1 protein fused to the C-terminus of the *P. hamadryas* APOL1 protein (the C-terminal 36 amino acids). The junction between human/gorilla and baboon occurs at the C-terminal end of the 4^th^ putative membrane spanning region, such that all chimeras encode the baboon C-terminal leucine zipper motif. (B) Kaplan-Meier survival curve of mice expressing chimeric APOL1 or human APOL1 by hydrodynamic gene delivery (HGD) inoculated with 5000 *T. b. rhodesiense* KETRI 243 parasites i.p. (*** p = 0.0001; Log-rank test). (C) Kaplan-Meier survival curve of WT and HET mice expressing chimeric APOL1 (Hum360) on a ubiquitin promoter, inoculated with 5000 *T. b. rhodesiense* KETRI 243 parasites i.p. (** p = 0.0035; Log-rank test). (D) Kaplan-Meier survival curve of mice expressing chimeric APOL1 (Gor360) or human APOL1 and HPR (ubiquitin promoters) by HGD inoculated with 5000 *T. vivax* IL1392 parasites i.p. (ns, p = 0.4302; Log-rank test). Experiment was repeated with a second isolate, with similar results.

### Transient expression of *APOL1* in mice does not protect from *T. vivax* infection

Using these optimized constructs, we interrogated the long-standing assumption that APOL1 also provides immunity to *T. vivax*^30^. The experiments were performed with uncharacterized field isolates of *T. vivax*, so we first confirmed the identity of the parasites by PCR using well-established *T. vivax*-specific primers (Supplementary Figure 5B). Upon species confirmation, we performed transient expression-based infections in mice producing either human APOL1 or the gorilla/baboon chimeric variant Gor360, and ultimately observed no APOL1-mediated protective effect in either group (Figure 7D and Supplementary Figure 5A). These data are consistent with studies using human serum administered to mice as a curative therapy, which revealed a general serum-resistance phenotype in *T. vivax* isolates^46^. This suggests that APOL1 alone, or in combination with HPR, is insufficient to provide protection against *T. vivax*.

## Discussion

Trypanosome lytic factors are primate-specific innate immune complexes that circulate in serum. The lytic component of TLFs is the APOL1 protein, which forms cation channels in membranes and leads to an osmotic swelling of *T. b. brucei* parasites. Here, we reveal that the baboon variant of the APOL1 protein is a broadly acting trypanocidal protein that is less restricted by pH for channel formation^8^. Further, we find that the baboon genus contains many variants of the APOL1 protein, and many of these variations have functional implications in protection against African trypanosomes. We have generated several genetically modified mice that express variations of the cDNA of one of these baboon APOL1 variants and have created a model system whereby the interaction between APOL1 and virtually any pathogen can be investigated. Since the *APOL1* expression system in our mice is not cell-type specific, our genetically modified mouse model is the best system that currently exists to analyze any APOL1-pathogen interactions in a physiological setting. As one example, our APOL1 transgenic mice are protected from *Leishmania major* infection^47^. *APOL1* expression is upregulated by interferon signaling, suggesting that it also plays an antiviral role, and indeed *APOL1* overexpression was shown to limit HIV replication *in vitro*^13^. We maintain that the study of primate APOL1 antimicrobial function and cell biology must involve the use of physiologically relevant systems, such as those presented here. In this manuscript, we are specifically interested in modelling the feasibility of genetically modified livestock resistant to trypanosome infection.

Before generating the mouse models of baboon APOL1 expression, we investigated the trypanolytic mechanism of baboon APOL1 *in vitro*. We have revealed that baboon APOL1 is less restricted by pH than human APOL1. Baboon APOL1 inserts at both acidic and neutral pH (human APOL1 - only acidic), and channels open at both acidic and neutral pH (human APOL1 - only neutral). This property of baboon APOL1 could explain the protein’s ability to lyse *T. b. gambiense* parasites, suggesting that APOL1 may insert into the endosomal membrane prior to reaching the TgsGP-containing compartments. However, under physiological conditions, the APOL1 protein will be delivered to trypanosomes by a TLF complex. Therefore, to insert in a lipid membrane, the APOL1 likely requires a yet-to-be-characterized TLF-disassociation step that may require a decrease in the environmental pH. These aspects of the mechanism cannot be studied using recombinant bacterially-derived proteins. The transgenic mouse models presented allow for the investigation of these mechanisms using human, baboon, or chimeric APOL1 proteins on HDLs as a TLF complex.

Recently evolved APOL1 variants in the human population are associated with glomerular diseases, where the source of damage is locally-produced APOL1 lysing the cells that produce it^48^. Our baboon APOL1 germline transgenic mice do not display any obvious negative phenotypes, while our chimeric baboon APOL1s (Hum360, heterozygote, and Gor360) had embryonic lethal phenotypes most likely driven by increased APOL1 expression during early development. In the context of modelling genetically engineered livestock, a more successful mouse model would express high enough concentrations of APOL1 to protect against all relevant African trypanosome species, but not at a level that causes toxicity to the animal. The variety of mouse lines presented here will offer excellent tools for the study of a variety of microbial infections, including trypanosomes.

APOL1 resistance in *T. congolense* and *T. vivax* has been observed in field isolates^49^ and *T. congolense* resistance can be induced by serial passage in the presence of low concentrations of human serum ^40^. We and others hypothesize that many livestock-infective trypanosomes are capable of developing APOL1 resistance, but it is also clear that these isolates rarely establish human infections with only approximately 20 documented cases of ‘atypical’ trypanosomiasis in the past century, only a small fraction of which were caused by *T. congolense*^50,51^. We hypothesize that while TLF mediates primate immunity to *T. b. brucei* and *T. b. evansi* infection, there are likely more factors involved that are responsible for cooperating with TLF to control *T. congolense*, *T. vivax*, *T. lewisi*^52^, *T. musculi*^53^, and other non-human infective, yet seemingly APOL1-resistant, parasites. One possible explanation involves the concentration of TLF in blood versus peripheral tissues and the fact that some trypanosomes are hypothesized to be preferentially blood-resident organisms. The low-TLF niches may select for the emergence of TLF-resistance in surviving parasites. These parasites, however, may have adapted an unfavorable pathogenesis mechanism to maintain this fitness as a result of being forced out of the bloodstream. These parasites, while TLF resistant, may then be incapable of establishing clinical infections in their primate hosts. More generally, it is possible that TLF resistance in *T. congolense* and *T. vivax* is not sufficient to mediate primate infectivity. These hypotheses could be addressed in the future by performing *in vivo* baboon challenges with TLF-resistant strains. These or similar studies seem particularly relevant given the maintained assumption that human immunity to *T. vivax* is mediated by APOL1^30^. The data presented here challenges that assumption, suggesting the mechanism of primate immunity to *T. vivax* and *T. congolense* is more complex than is currently understood. APOL1, if it does play a role in immunity to *T. vivax*, is clearly not sufficient alone (or when co-expressed with HPR). Notably, there remains debate with respect to whether data obtained from *T. vivax* experiments performed in mouse models can be clearly interpreted. Given that the majority of *T. vivax* isolates cannot be cultivated in mice, this calls into question the relevance of those isolates that can infect mice^30^.

In order to create animal models that are protected against trypanosome infections as well as primates are, it stands to reason that the APOL1 and HPR proteins would need to co-localize onto the same HDL particles, thus recreating TLFs. So far, it has proven difficult to achieve this in transgenic mice. We are unsure of the cause; there may be certain HDL proteins and other HDL synthesis mechanisms that play roles in primate TLF assembly that have not yet been characterized and do not exist in mice. TLF assembly has generally not been extensively investigated, although we suggest that expanding this literature may prove useful in the contexts of APOL1-related immunology as well as APOL1-related tissue toxicity. Nevertheless, a second strategy for generating fully protected animal models is to increase APOL1 protein expression level. The various genetic constructs generated in this manuscript were created with both of these challenges in mind, and at least the latter challenge has been met with our chimeric APOL1 proteins. The chimeric protein strategy may be the more effective option than co-expression of APOL1 and HPR in the context of genetically engineered livestock because some veterinary trypanosomes express HpHbR receptor variants that do not bind HPR.

It remains unclear why there is less baboon APOL1 in serum than the human protein. Given that the models presented here used the same expression system with either baboon or chimeric protein, and had the same levels as the baboon and human, we hypothesize that this differential concentration is not based on RNA expression. We instead suggest that the baboon APOL1 protein is not as efficiently secreted and loaded onto HDLs, or that baboon APOL1 is degraded by serum proteases more efficiently than human APOL1 after secretion. The C-terminal region of baboon APOL1, which is necessary and sufficient for *T. b. rhodesiense* resistance does not apparently decrease the efficiency of HDL loading or protein secretion as evidenced by the chimeric APOL1 proteins created here (Figure 7). However, these chimeric proteins were evidently produced too efficiently, given the associated developmental fitness cost. We believe that the chimeras are strong candidate genes to be used in future murine and livestock studies after titrating their expression levels with slightly weaker promoter systems.

In summary, we have shown that APOL1 proteins are potent ion channel-forming proteins that can be used in genetically modified models to create disease-resistant animals. Our long-term research goal is to investigate the possibility of creating trypanosome-resistant livestock that could thrive in the tsetse endemic regions of sub-Saharan Africa through transgenic expression of *APOL1*. To this end, we have recently used somatic cell nuclear transfer to produce the first cloned Kenyan Boran bull in Africa^54^. Trypanosome-resistant cattle breeds could expedite agricultural development and significantly reduce the rates of human trypanosomiasis transmission in sub-Saharan Africa^55^, especially when combining their impact with additional control efforts. However, the fact remains that if APOL1 does not provide protection against *T. vivax* infection, then these perspectives may need to be reconsidered or restricted to *T. vivax*-free areas of Africa. Finally, we hope that the biology presented here may assist us and others in our efforts to illuminate the evolutionary relationship between primate *APOL1* and trypanosomes.

## Methods

### Parasites

*T. b. brucei* Lister 427 and *T. b. gambiense* ELIANE were grown *in vitro* in HMI-9 with 10% fetal bovine serum and 10% Serum Plus at 37°C in 5% CO_2_. *T. b. brucei* Lister 427-SRA is a human APOL1 resistant line generated by the Cross lab that expresses the SRA gene from *T. b. rhodesiense*. The other trypanosomes were maintained exclusively in mice: *T. b. rhodesiense* KETRI 243, *T. b. brucei* AnTaT1.1 DRF, *T. b. evansi* Antat 3, *T. vivax* IL1392, and *T. congolense* STIB 68-Q.

### *T. vivax* species confirmation

DNA was isolated from *T. vivax*-infected mouse blood or uninfected mouse blood. DNA was amplified using established thermocycling conditions and *T. vivax*-specific primers ^56^. Forward: GCCATCGCCAAGTACCTCGCCGA Reverse: TTAAAGCTTCCACGAGTTCTTGATGATCCAGTA

### Primate samples

Primate plasma samples were obtained from the Southwest National Primate Research Center at the Texas Biomedical Research Institute.

### Antibodies

For the detection of baboon APOL1, we generated a novel antibody through AnaSpec Incorporated, which was raised in rabbits against the following peptide: CSVEERARVVEMERVAESRTTEVIRGAKIVDK. An anti-human APOL1 antibody (ProteinTech, 11486-2-AP) was used for the detection of human, gorilla, or the chimeric APOL1. The anti-human HP that recognizes baboon HPR is commercially available (Sigma, H8636). The secondary antibody used in all western blots was the anti-rabbit TrueBlot conjugated to HRP (1:5000) (Rockland Antibodies, 18-8816-33).

### Phylogenetic analysis of baboon *APOL1*

The genotype data generated originally generated by the Baboon Genome Project were downloaded at ftp://ftp.hgsc.bcm.edu/Baboon/Panu_2.0/ which is contributed by Baylor College of Medicine Human Genome Sequencing Center (HGSC). The data set consists of 16 individuals belonging to six species within the genus *Papio*, and *Theropithecus gelada*, a member of a closely related genus that serves as an outgroup^29^. We extracted variant calls in APOL1 exons by filtering nucleotide position using VCFtools (version 0.1.17)^57^, where we retrieved 39 single nucleotide polymorphism (SNP) sites and 3 indels. The variant sites of the 32 samples were aligned using a customized PERL script and a maximum likelihood tree of the haplotypes was inferred with FastTree (version 2.1.8)^58^. The haplotypes and the tree were then visualized in Adobe Illustrator 2018. To estimate nonsynonymous (dN) and synonymous (dS) substitution rates and thereby detect adaptive selection on APOL1 gene, we reconstituted the full-length codon sequences (1167 nt) of the 32 haplotypes according to a reference sequence. The reference sequence, consisting of 1221 nucleotides, is the mRNA of APOL1 in *Papio anubis* (NCBI accession NM_001302101). We matched the SNP sites to the reference sequence position by position using a customized pipeline. The dN/dS ratio was then estimated using program PAML V4.9^59^ with a neutral model (M0), a nearly neutral model (M1a) and a positive selection model (M2a). Likelihood ratio tests were carried out between models. **Production and purification of rAPOL1s.** We expressed and purified N-terminally 6xHIS-tagged full length rAPOL1 using the pNIC vector from *E. coli* BL21 Codon Plus RIPL cells (Agilent) grown in Overnight Express media (Novagen) using a previously described procedure with slight modification. Inclusion bodies were washed in the presence of the protease inhibitors AEBSF and EDTA, then solubilized in 1% zwittergent 3-14 (SB3-14, EMD-Millipore). Solubilization was facilitated by briefly adjusting to pH 12 with 10 mM NaOH and 150 mM NaCl for one minute before adding 30 mM Tris-HCl (pH 7.4) to reduce the pH to 8.0. APOL1 proteins were then separated on a Superdex-200 16/60 size-exclusion column (GE Life Sciences) equilibrated in 50 mM Tris-HCl, pH 8.5, 150 mM NaCl, and 0.5% SB3-14. APOL1-containing fractions were pooled and bound to a nickel affinity column (HisTRAP, GE Life Sciences) equilibrated in 20 mM Tris-HCl, pH 8.5, 150 mM NaCl, 10 mM imidazole and 0.5% SB3-14 and eluted with a 10-500 mM imidazole gradient in the same buffer. APOL1 proteins were purified from any remaining detectable contaminants via a second size-exclusion protocol using the same size-exclusion column equilibrated in 50 mM Tris-HCl, pH 8.5, 150 mM NaCl, and 0.05% n-Dodecyl-beta-Maltoside (DDM) (Thermo Scientific). Purified proteins were flash-frozen and stored at −80°C.

### Western blotting

All western blots were performed using PVDF membranes after overnight electro transfer at 30 mV. Membranes were blocked with 5% milk powder in 150 mM NaCl, 50 mM Tris, pH 7.5. Membranes were probed with both primary and secondary antibodies diluted in the blocking buffer with interspaced washes (3x 10 minutes) with 150 mM NaCl, 50 mM Tris, pH 7.5 prior to chemiluminescent visualization (Thermo Scientific). All washes and probes were conducted in the presence of 0.05% Tween-20 (Fisher Scientific).

### Trypanosome *in vitro* lysis assays

Cultured *T. b. brucei* Lister-427 parasites were diluted to 5 x 10^5^ cells/ml in in culture media as described above and 100 μL was added to each well of an opaque 96 well plate. Parasites were then diluted 1:1 with various concentrations of rAPOL1 proteins suspended in HMI-9. The concentration of DDM in the assay was maintained below the theoretical critical micelle concentration (0.006%) and did not affect parasite viability. After 20 hours of incubation at 37°C with 5% CO_2_, 20 μL of alamarBlue (Invitrogen) was added to each well. The assay involves the reduction of resazurin by metabolically active cells to the fluorescent resorufin, allowing quantitation of non-lysed cells at 4 hours post-reagent addition in a spectrofluorometer.

### *In vivo* parasite infections

All experiments conducted in mice were approved by the IACUC committees of the appropriate institutions. For trypanosome infections, a total of 5000 parasites (unless otherwise indicated) was injected intraperitoneally (i.p.) into transgenic or wild type C57BL/6N-J mice derived from founder mice expressing baboon APOL1, baboon HPR, and/or human APOA-I after breeding with both the C57BL/6N and J strains of wild type mice. Genotype was confirmed by PCR to determine heterozygote vs homozygote status of each individual transgene. Parasitemia was monitored by tail bleeding. Mice were euthanized by isoflurane when parasitemia reached 1 x 10^9^ parasites/mL. Mice that were transiently transfected with any HGD construct were infected 48 hours after plasmid injection. All survival experiments were analyzed statistically using the log-rank test via GraphPad Prism. The *T. vivax* infections were performed at the International Livestock Research Institute in Nairobi, Kenya. These studies used Swiss Webster mice and were repeated with two different *T. vivax* isolates with similar results.

### Lipoprotein isolations

The density of baboon or mouse plasma was adjusted to 1.25 g/ml using KBr and samples were ultracentrifuged in an NVTi-65 rotor (Beckman Coulter) for 16 hours at 49,000 RPM, 10°C. The top 33% of the resulting gradient was aspirated and dialyzed against 5 changes of 150 mM NaCl, 50 mM Tris, 0.5 mM EDTA, pH 7.5 before storing at −80°C.

### De-glycosylation of baboon lipoproteins

PNGase F was obtained from New England BioLabs and de-glycosylation was performed in 50 μL under denaturing conditions according to the manufacturer’s instructions except that 1 mM EDTA, 1 mM EGTA, 1 mM AEBSF (final concentrations), and 0.33 μL of the HALT protease inhibitor cocktail (Thermo Scientific) were added to the lipoproteins prior to denaturation.

### Generation of transiently transgenic mice

Expression of baboon *HPR*, or chimeric *APOL1s*, was achieved in certain cases using hydrodynamics-based gene delivery (HGD)^18^. Briefly, mice were injected through the tail vein with 10% of their corresponding body weight in sterile 0.9% NaCl solution that contained 25 μg of plasmid DNA encoding the cDNA of the gene of interest flanked by an SV40 polyadenylation sequence and a beta-globin intron under the control of a human ubiquitin promoter. Prior to trypanosome infections, blood was drawn from the tail two days post-transfection to confirm protein expression by western blot (Supplementary Figure 4).

### Histology

Kidneys were excised from >1 year-old mice. The kidneys were sliced into two longitudinal halves and were washed in phosphate buffered saline before being fixed overnight in 4% paraformaldehyde at 4°C. The kidneys were then transferred to 70% ethanol in phosphate buffered saline for storage. The kidneys were submitted to the New York University pathology core facility. 5 μm sections were cut, fixed in formalin, and embedded in paraffin prior to being stained with hematoxylin and eosin.

### *T. congolense* quantitative PCR

To design primers for quantitative PCR specific to *T. congolense* parasites, we targeted the *T. congolense cathepsin L* (*CATL*) gene. *CATL* is a multicopy gene, existing in up to 20 copies in some isolates, making it an ideal candidate gene for highly sensitive PCR reactions. Using previous published broadly specific *CATL* primers, we amplified and subsequently sequenced the gene from *T. congolense* STIB 68-Q. We then designed primers (F: 5’-GATCTTCGCACCAACGACCT, R: 5’-AAGGGCAATGTGTTCACGGA) that amplified a 90 bp segment of the gene for quantitative PCR. The quantitative PCR cycling conditions are as follows: Initial denaturation at 94°C for 1 minute, followed by 38 cycles of 98°C for 5 seconds, 59.5°C for 30 seconds, and 72°C for 20 seconds. PCRs were performed in 20 μL final volumes consisting of 10 μL of SYBR select Master Mix (Applied Biosystems), and 10 μL of remaining components amounting to 30 ng of total template DNA and 0.5 μm primer concentrations. **Tissue collection for quantitative PCR.** Infected mice were manually transcardially perfused through the left ventricle after aspirating the right atrium with 20 ml of ice-cold heparinized phosphate buffered saline. After cutting the atrium and prior to starting the injection of saline, 100 μL of blood was collected from the thoracic cavity and immediately pipetted into 500 μL of TriReagent (Zymo Research). After perfusing the animal, sections of the lung, liver, gonadal adipose depot, spleen, and brain were excised, washed in phosphate buffered saline to remove residual blood, weighed, and placed into 500 μL of TriReagent (Zymo Research). Tissue samples were stored at 4°C until homogenization by sequential needle passing. Tissue homogenates were then mixed with an equal volume of molecular grade ethanol before isolating genomic DNA using a commercially available kit (DNeasy Blood and Tissue Kit - Qiagen).

### Electrophysiology

Planar lipid bilayers were formed at room temperature from soybean asolectin [lecithin type IIS (Sigma Chemical) from which nonpolar lipids had been removed] and cholesterol (Sigma, C8667) across an 80- to 120-μm hole in a Teflon partition separating two solutions of bilayer buffer (1 mL vol), as described previously^9^. Bilayer buffer: 150 mM KCl, 5 mM CaCl_2,_ 0.5 mM EDTA, 5 mM K-succinate, 5 mM K-HEPES, pH 5.5-7.5. Briefly, 20 μL of lipid (1.5% asolectin 0.5% cholesterol) in pentane was layered on top of the solutions and the pentane was allowed to evaporate. Bilayers were formed by raising each solution above the hole that was pretreated with squalene. Bilayer formation was monitored by measuring a change in capacitance. Voltages were maintained using the BC-535C bilayer clamp (Warner Instruments) and are given as the voltage of the *cis* solution (defined as the side to which protein was added) with respect to the *trans* solution. The current response was filtered at 30 Hz by a low-pass eight-pole Bessel filter (Warner Instruments) and recorded using IGOR NIDAQ Tools MX 1.0 and IGOR software (WaveMetrics) via an analog-to-digital converter (NI USB-6211; National Instruments).

### Generation of chimeric *APOL1* genes

The “Hum360” chimera was obtained from Thomson and Genovese *et al*.^20^. To generate the “Gor360” chimera, the gorilla (*Gorilla gorilla gorilla*) *APOL1* gene was first assembled through PCR of each individual *APOL1* exon from gorilla genomic DNA obtained from the Wildlife Conservation Society in Bronx, NY. The exon flanking primers were designed using a complete, assembled gorilla genome (gorGor_Susie3) obtained from the Eichler Lab at the University of Washington European Nucleotide Archive, Project Accession PRJEB10880, Taxonomy_ID: 9595^60^. Exon 1 F: TGGTCATGGAGGTCAGGATATCGAG; Exon 1 R: CTAGAAGAAGCCCAGATGGCCC. Exons 2 and 3 were sequenced simultaneously because the intron separating the two sequences is relatively small (182 bp). Exons 2 and 3 F: CAGGCCCTGGTCATTGTCAG; Exons 2 and 3 R: CTTGGGGCAGACTCATTGGC. Exon 4 F: GGCTGTTATGCACTCCCAC; Exon 4 R: CTGCCTGGAGGAGGTGTG. Exon 5, the largest exon, was sequenced via a nested PCR. Exon 5 nest F: CTGCCTGGAGGAGGTGTG; Exon 5 nest R: GCATTTTGTCCTGGCCCCG. Exon 5 inner F: GCATTTCCTCTGGCATCCTGAC; Exon 5 inner R: CACGGAGCCTTCTTATGTTA. The sequenced exons were stitched together manually *in silico*, and the full-length gene was then synthesized by Invitrogen GeneArt Gene Synthesis Service by Thermo Fisher Scientific. The “Gor360” chimera was then created through In-Fusion cloning (Takara) by fusing the first 1080 bp of the gorilla gene with the C-terminal region of *P. hamadryas APOL1* (accession: FJ429176). All PCRs were performed using Pfu Ultra AD high-fidelity polymerase (Agilent Technologies).

### Generation of targeted *P. hamadryas APOL1*-expressing mice

For targeting constructs containing the *P. anubis APOL1* genomic sequence, all sequences were isolated from a BAC obtained from the Children’s Hospital Oakland Research Institute. All genomic coordinates are from the Panu_3.0/papAnu4 version of the *P. anubis* genome in the UCSC genome browser. First, a floxed neomycin expression cassette was inserted into the first intron of *APOL1*, which removed 40bp from the intron (chr10:77,080,500-540). The modified *APOL1* gene (chr10:77,075,579-77,097,083) containing the neomycin (neo) selection cassette was transferred to a R6K origin of replication plasmid. For the Rosa26 targeting vector, a splice acceptor and polyadenylation sequence from the rabbit beta globin gene was inserted upstream of the *APOL1* promoter to reduce readthrough transcription from the Rosa locus promoter. The two targeting vectors were then constructed by using homologous recombination in *E. coli* to insert the transgene sequences from the R6K constructs into BAC vectors containing either mouse genomic sequences from the Rosa26 safe harbor or the Myh9 loci, as described^61^. The *P. anubis* APOL1-Myh9 construct was converted to *P. hamadryas APOL1* using homologous recombination to insert the necessary point mutations.

For the UBC-APOL1 construct, the vector was first assembled in an R6K plasmid and consists of the human ubiquitin C (UBC) promoter and the rabbit beta globin intron upstream of a *P. hamadryas* APOL1 cDNA, followed by the bovine growth hormone polyadenylation sequence. For selection in embryonic stem cells, a self-deleting neomycin resistance cassette was placed downstream of the polyadenylation site. For the Alb-HPR-UBC-APOL1 construct, a fusion of the mouse albumin enhancer and promoter sequences ^62^ and a baboon HPR cDNA with a beta globin polyadenylation sequence was inserted upstream of the UBC promoter in the UBC-APOL1 construct. For the UBC-APOL1-UBC-HPR construct, a UBC-driven HPR was inserted downstream of the UBC-driven APOL1 cDNA, and both cDNAs used an SV40 polyadenylation sequence. Targeting vectors were constructed by using homologous recombination in *E. coli* to insert the transgene sequences from the R6K constructs into BAC vectors containing mouse genomic sequences from the Rosa26 locus, as described above.

Constructs for the chimeric APOL1 mice (Hum360 *APOL1* and Gor360 *APOL1*) were fully synthesized (GenScript) in an R6K vector. Synthesized DNA included the UBC promoter, the rabbit beta globin intron, the Hum360 or the Gor360 chimeric *APOL1* coding sequence, SV40 polyadenylation sequence and followed by a self-deleting neo resistance cassette. Bacterial homologous recombination was used to insert the construct into a BAC vector for targeting the Rosa26 locus. All vector sequences are available upon request.

All constructs were electroporated into either C57BL/6NTac or hybrid C57BL/6NTac:129S6/SvEvTac ES cells, and correct targeting of the two loci in mouse ES cells was confirmed using loss- and gain-of-allele PCR analysis. All antibiotic selection cassettes were removed using either a Cre recombinase expression vector in the targeted ESC clone or during male germline maturation. Heterozygous targeted cells were microinjected into 8-cell embryos from Swiss Webster albino mice (Charles River Laboratories), yielding F0 VelociMice that were 100% derived from the targeted cells^63^. These mice were subsequently bred to homozygosity and maintained in the Hunter College Animal Facility during the study period. All relevant protocols were approved by the corresponding Institutional Animal Care and Use Committees of Regeneron and Hunter College.

## Acknowledgements

We thank the Baboon Genome Consortium, specifically Dr. Jeffery Rogers and Dr. Muthuswamy Raveendran, for early access to the baboon genome sequences. We thank the New York University Langone Experimental Pathology research facility for processing the mouse kidney sections. We thank Matthew Koss at Regeneron for conducting all mouse genotyping. The authors declare no conflicts of interest. Funding for this project was graciously provided by: National Science Foundation IOS-1249166

## Abbreviations used

APOA-I: Apolipoprotein A-I
APOL1: Apolipoprotein L-1
CATL: Cathepsin L
G0/G1/G2: Genotype 0/1/2
HDL: High-density lipoprotein
HET: Heterozygote;
HGD: Hydrodynamic gene delivery
HOM: Homozygote
HpHbR: Haptoglobin-hemoglobin receptor
HPR: Haptoglobin-related protein
I: Current
I.p.: Intraperitoneal injection
PNGase F: Peptide N-glycosidase F
TLF: Trypanosome Lytic Factor
V: Voltage.

## Author Contributions

JV, SF, KL, SP, DBK, DK, BG, AR, LL, WQ, and RT conducted the experiments in the manuscript and conceived the hypotheses along with JR. Specifically, JV performed all gels with SF and DBK, protein purifications and in vitro parasite assays; JV, DK, SF and KL conducted mouse infection experiments; SP designed and conducted the qPCR-based experiments with JV; BG and AR cloned the gorilla APOL1 gene; LL and WQ performed the phylogenetic analyses; and RT and KL performed the electrophysiological experiments. CS, JSR and AE designed and generated the murine targeting plasmids and transgenic mice. JV, SF, and JR wrote the paper.

---

**Supplementary Figure 1.**
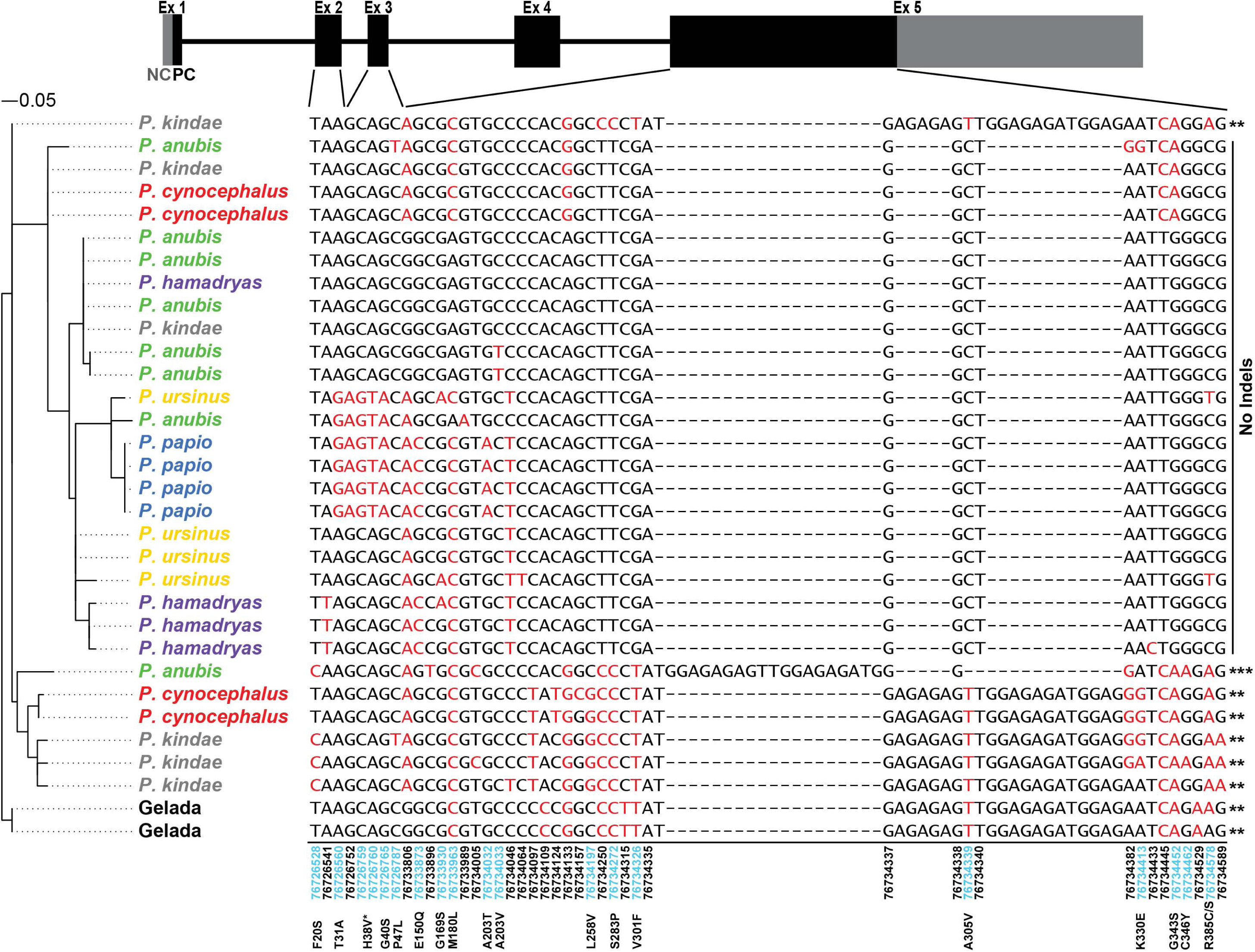
Phylogenetic analysis of the baboon *APOL1* gene. Pictured above the phylogenetic tree and haplotype table is a schematic of the five protein-coding exons of *APOL1* in primates (NC, non-coding; PC, protein-coding). All of the observed polymorphisms were present in exons 2, 3, and 5. Only the sites that are polymorphic between the baboon individuals are shown. Each observed haplotype is depicted separately, with haplotypes grouped by their phylogeny (left). Polymorphisms that do not match the reference sequence (Panu2.0) are red, while reference-matching sites are black. The genetic coordinates of each site in the reference sequence are listed below the table, and polymorphic sites that encode missense mutations (relative to the reference sequence Panu 2.0) are highlighted by the encoded amino acid change above the table. H38V (*) is intentionally located between two SNPs because the amino acid change is encoded by two adjacent polymorphisms. The ‘No Indels’ and asterisks at the right refer to supplementary table 2, wherein the amino acid changes, starting at amino acid position 300, encoded by the different Indel patterns are clarified.

**Supplementary Figure 2.**
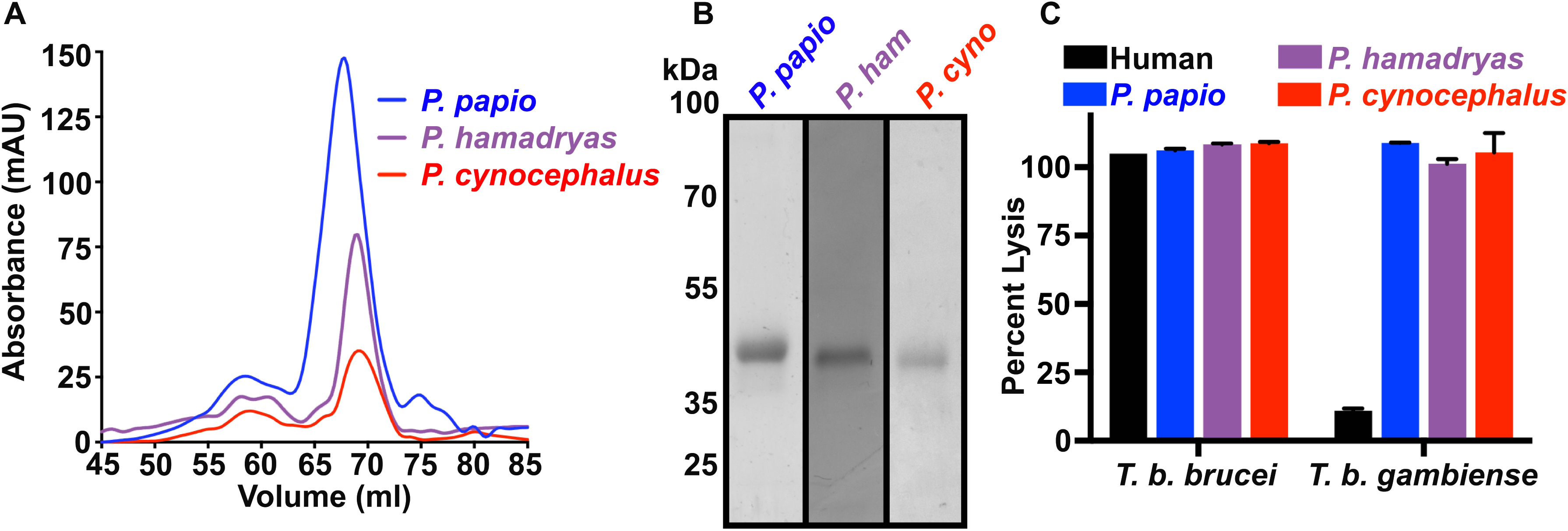
Purification of baboon APOL1 made recombinantly in bacteria. (A) UV absorbance chromatograms of the final size exclusion chromatography step of three different baboon recombinant APOL1 (rAPOL1) protein purifications. (B) Coomassie stained gels of the purified *Papio* rAPOL1 proteins. Expected size: 42 kDa (C) 24-hour trypanolysis assay showing the lytic capacity of 1,250 ng/ml of each of the indicated rAPOL1 proteins against *T. b. brucei* and *T. b. gambiense*. Reproducible data from multiple assays were combined to generate this graph (error bars represent mean +/− SD of three experimental replicates).

**Supplementary Figure 3.**
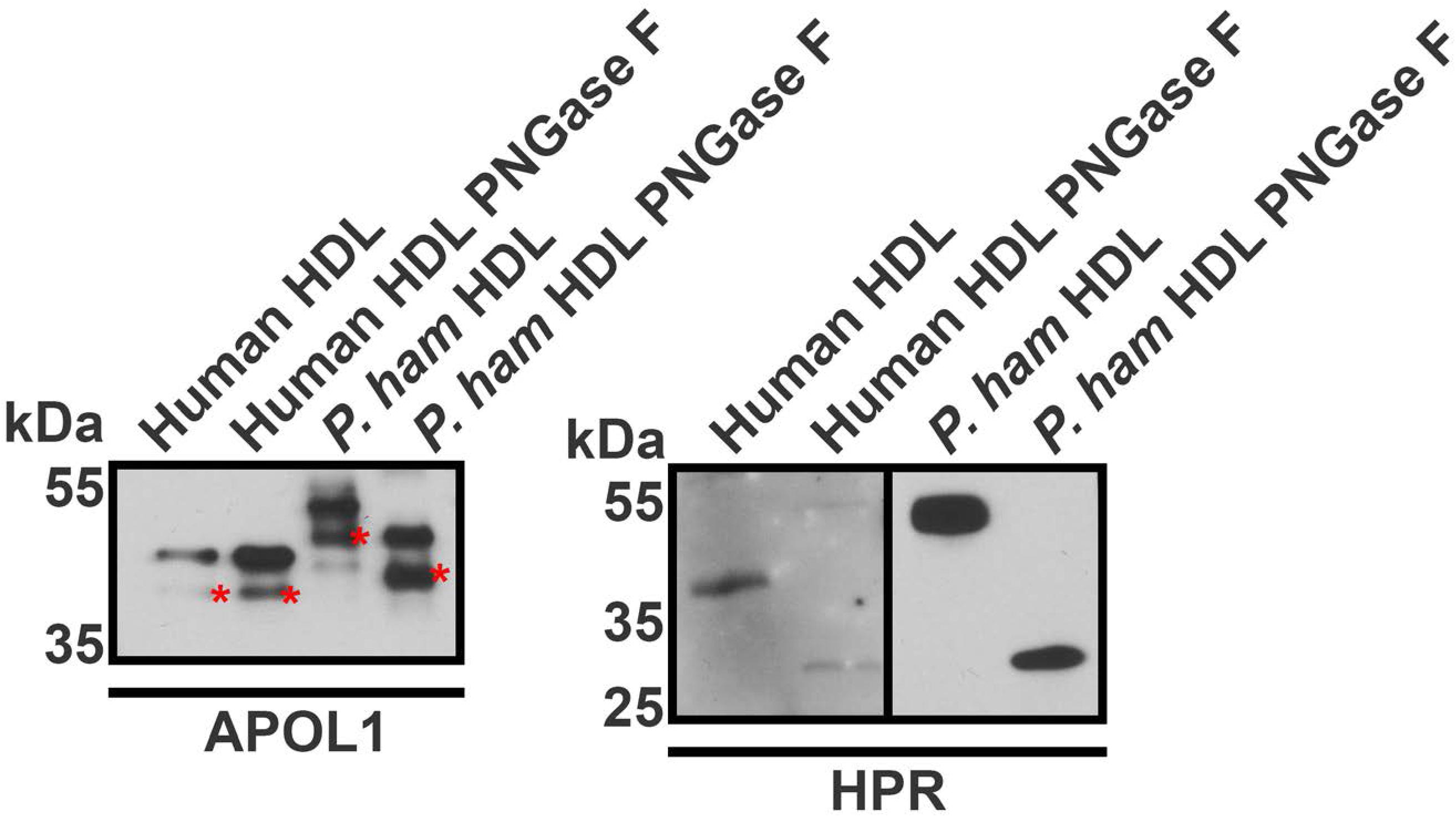
Human and Baboon HPR and APOL1 proteins are post-translationally modified. Total HDL lipoproteins were isolated from human and baboon plasma. The HDLs were either treated with vehicle control or PNGase F before electrophoresis under reducing conditions. The molecular mass of baboon APOL1 and both human and baboon HPR decrease significantly after PNGase F treatment, indicating that the proteins are *N*-glycosylated. The red asterisk in the left panel highlights an APOL1 C-terminal cleavage product that often accumulates during sample collection.

**Supplementary Figure 4.**
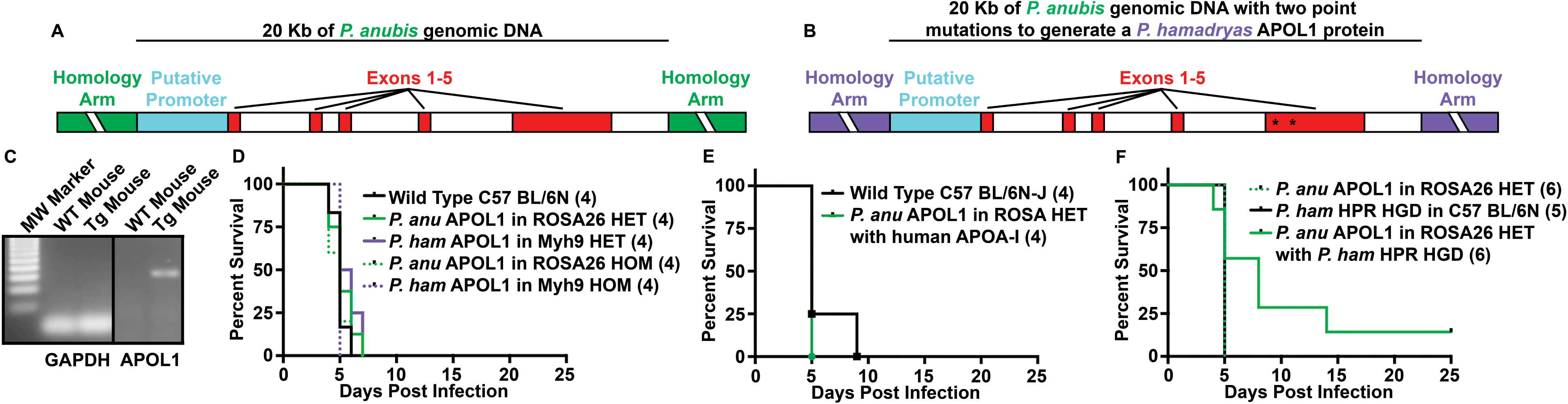
Integration of the genomic baboon *APOL1* locus in mice is not sufficient to protect from *T. b. brucei*. (A-B) Schematics of the targeting constructs used to insert the genomic baboon *APOL1* locus into the mouse genome. The construct in panel B was generated by making two substitutions (*) in the construct shown in panel A (E150Q and L180M). (C) Ethidium bromide-stained agarose gel showing the results of gene-specific PCRs of cDNA synthesized from RNA derived from liver homogenates of a wild-type mouse and a germline transgenic mouse expressing *P. anubis APOL1* from the *ROSA26* locus. (D-F) Kaplan-Meier survival curves of mice infected with 5000 *T. b. brucei* (427-SRA) parasites intraperitoneally (i.p.). (D) Heterozygous or homozygous transgenic mice expressing either *P. hamadryas APOL1* from the Myh9 locus or *P. anubis APOL1* from the ROSA26 locus compared to wild-type counterparts. (E) *P. anubis APOL1* and human *APOA-I-*expressing mice compared to wild-type counterparts. (F) *P. anubis APOL1*-expressing mice expressing *P. hamadryas HPR* by HGD compared to *APOL1* or *HPR* alone (p = 0.09; Log-rank test).

**Supplementary Figure 5.**
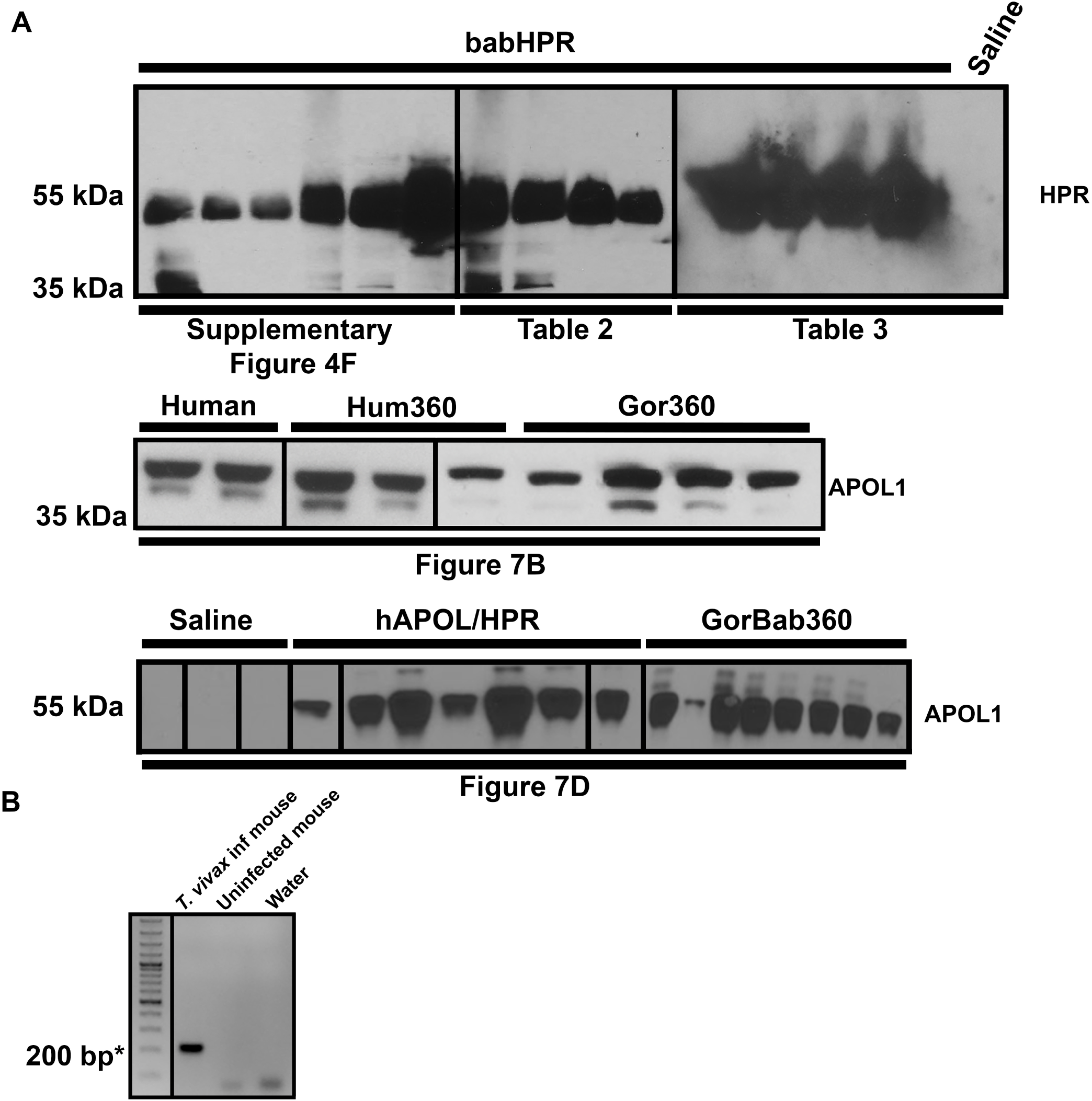
Validation of protein expression and parasite identity in transiently transgenic mice (hydrodynamic gene delivery). (A) The mice expressing either *HPR,* or APOL1 and HPR, or the *APOL1* chimeras via HGD in various experiments were assayed for protein production by western blot of 0.75 µL of plasma collected 2 days-post vector injection by HGD (the same day as parasite infection) to confirm that the data could be interpreted without consideration of poor protein expression. (B) *Trypanosoma vivax* (IL1392) identity was confirmed by PCR from DNA extracted from blood of infected and uninfected mice, using established primers. The expected band size was 200 bp, indicated with an asterisk.

**Westerns.**
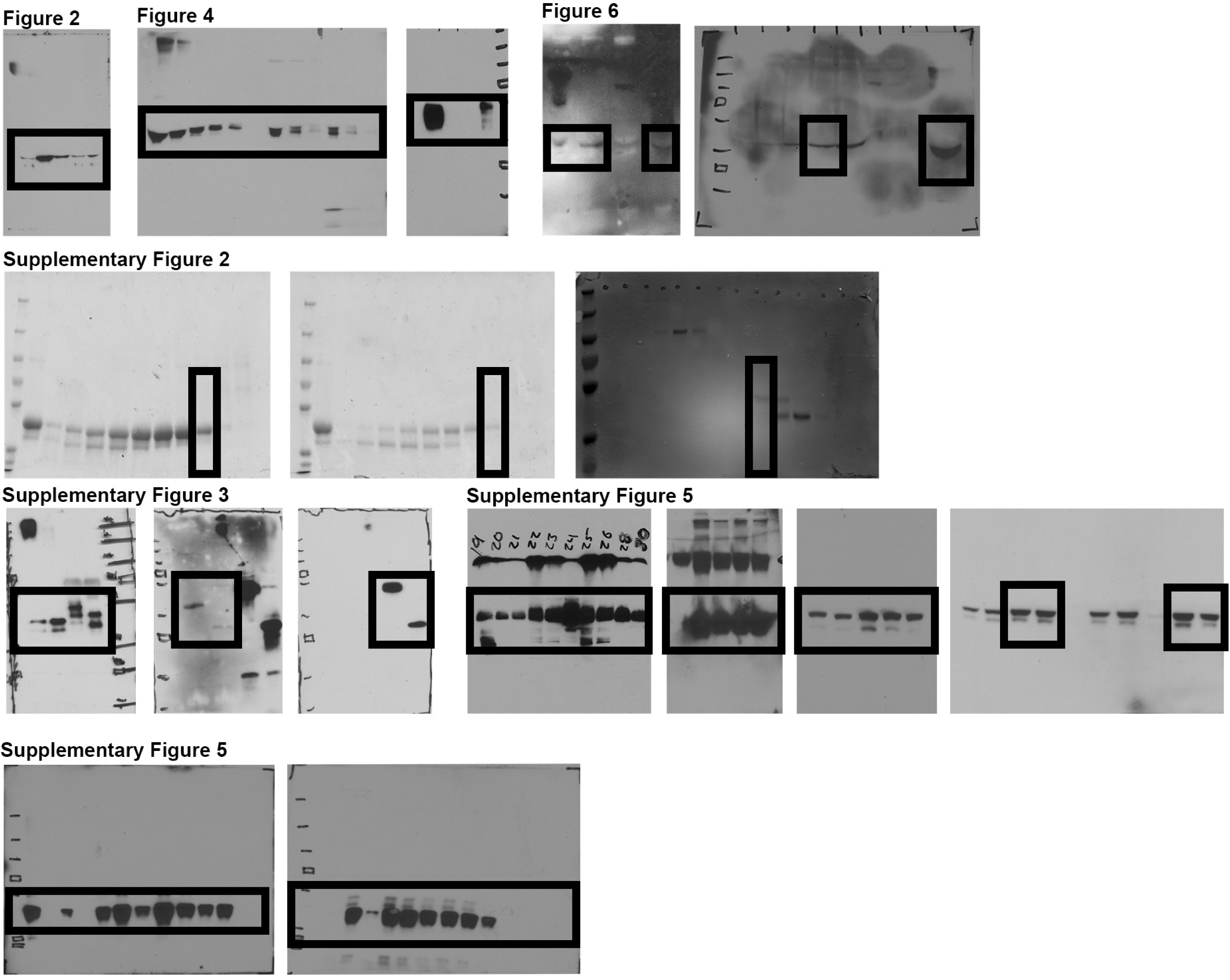
All raw western blots and Coomassie stains in this manuscript.

**Supplementary Table 1.** Sources of *Papio* DNA used in this study.

| Sample ID | Species | Biomaterial Provider Institution/Wild | DNA Source | Sex |
| --- | --- | --- | --- | --- |
| L142 | Papio anubis | Southwest National Primate Research Center (SNPRC) | WBC | M |
| LIV5 | Papio anubis | Southwest National Primate Research Center (SNPRC) | Liver | F |
| 28697 | Papio ursinus | Southwest National Primate Research Center (SNPRC) | N/A | F |
| 28755 | Papio ursinus | Southwest National Primate Research Center (SNPRC) | N/A | F |
| 28547 | Papio papio | Southwest National Primate Research Center (SNPRC) | N/A | N/A |
| 30388 | Papio papio | Southwest National Primate Research Center (SNPRC) | N/A | M |
| 34474 | Papio kindae | BCM/Blood collected in the wild - in southwest Zambia | Blood | F |
| 34472 | Papio kindae | BCM/Blood collected in the wild - in southwest Zambia | Blood | M |
| 34449 | Papio kindae | BCM/Blood collected in the wild - in southwest Zambia | Blood | F |
| 30877 | Papio anubis | Yerkes National Primate Research Center (YNPRC) -<br>Aberdare region of Kenya | WBC | N/A |
| 30977 | Papio anubis | Yerkes National Primate Research Center (YNPRC) -<br>Aberdare region of Kenya | WBC | N/A |
| 16066 | Papio cynocephalus | Baylor College of Medicine - collected in central Tanzania | WBC | F |
| 16098 | Papio cynocephalus | Baylor College of Medicine - collected in central Tanzania | WBC | F |
| 97074 | Papio hamadryas | Washington University School of Medicine and NYU -<br>Awash Park, Ehtiopia | WBC | F |
| 97124 | Papio hamadryas | Washington University School of Medicine and NYU -<br>Awash Park, Ethiopia | WBC | M |

**Supplementary Table 2.**
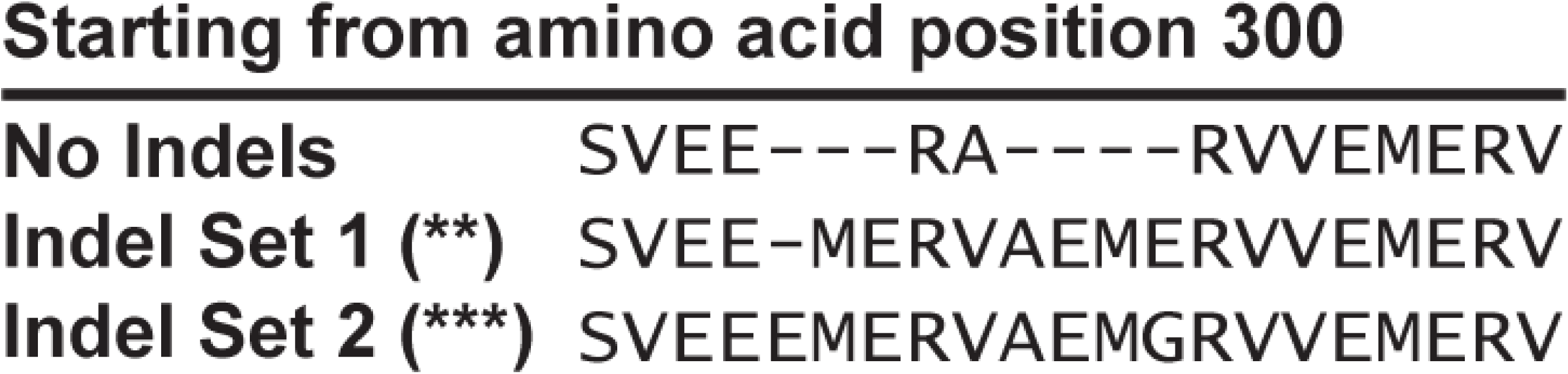
Amino acid sequences of *Papio* APOL1 proteins generated by the observed Indels. Asterisks correspond to those present in Supplementary Figure 1.

